# Fine-scale survey of intertidal macroalgae reveals recent changes in a cold-water biogeographic stronghold

**DOI:** 10.1101/2022.02.25.482036

**Authors:** Cátia Monteiro, Joana Pereira, Rui Seabra, Fernando P. Lima

**Author notes:** These authors have contributed equally to this work and share first authorship (Cátia Monteiro, Joana Pereira). **Correspondence** Fernando P. Lima.

## Abstract

Global warming has been causing severe impacts on marine ecosystems, a notorious one being shifts in the geographical ranges of species. The north-western coast of the Iberian Peninsula is an especially interesting zone to study distributional shifts as it has a strong latitudinal thermal gradient, is influenced by the Canary upwelling system (which partially cancels coastal warming) and holds some of the most diverse macroalgae communities in Europe. Notably, it is within this region that many cold-water species, common in northern Europe, have their southernmost distribution refuge. Recent studies hypothesize that the environmental conditions may be nonetheless changing and already threatening this biodiversity hotspot.

The main goal of this study was to carry out a fine-scale assessment of the distributional limits of several macroalgae in North-western Iberia, as well as identify possible population and range shifts using historical data (2001-2005) as reference. In addition, invasive species were also surveyed. We also assessed if the regions of (i) Galicia, (ii) Northern Portugal, and (iii) Central Portugal displayed distinctive characters regarding macroalgae composition and abundance.

We identified an increase in abundance of some invasive macroalgae as well as a decrease in the abundance of some cold-water species. In the most severe cases, cold-water species were extirpated along hundreds of km. The compounded effect of the decrease in the abundance of cold-water species and the increase in the abundance of invasive species is leading to the homogenization of macroalgae communities in north-western Iberia.

## 1 Introduction

Climate change is affecting the biodiversity, structure and function of coastal ecosystems, especially in rocky intertidal zones (Jurgens et al., 2015; Poloczanska et al., 2016; Ullah et al., 2018). One of the most pervasive effects of climate change in marine species has been the change in their distributional ranges, with species of temperate or boreal origin disappearing from sites where temperatures surpass their higher thermal tolerance limits, and with tropical or warm-temperate species invading locations once unfavorably cold. While the first is concerning because we risk losing biodiversity and/or genetic diversity with cascading effects throughout the ecosystem, the second could also be damaging, especially in the case of invasive species, owing to their characteristic potential to alter entire communities (Stachowicz et al., 2002; Hawkins et al., 2009; Antão et al., 2020).

Near the distribution limit of a species, populations are expected to experience sub-optimal conditions that might render them more vulnerable than populations in the center of the distribution (Hampe and Petit, 2005). This leads to greater impacts and lower recovery capacity of these populations in the case of changes at large spatial scales, such as those caused by climate change (Pearson et al., 2009; Viana et al., 2014; Rubal et al., 2015). Macroalgae play a pivotal role in coastal ecosystems, given that they are primary producers, provide habitat and nursery grounds for a wide variety of species (of which many are economically relevant), and their canopies offer a buffer from the harsh physical conditions of the intertidal (Smale et al., 2013; Seitz et al., 2014; Bulleri et al., 2018). Crucially, macroalgae are sensitive to changes in temperature, which is an important driver of their key life cycle events, such as reproduction and growth (Harley et al., 2012). Therefore, it is important to study marginal populations of cold-adapted macroalgae, as they have a higher probability of disappearing first in response to climate change (Martínez et al., 2015), giving some insights into what may happen to other populations in the future.

In parallel, invasive species are spreading at an ever-faster pace due to the increase in vectors of introduction (mainly maritime traffic and marine aquaculture) and the facilitation from climate change (McKnight et al., 2021). Monitoring biological invasions is necessary as invasive species are at the forefront of ecological change by outcompeting native taxa or acting as vectors or agents of diseases to which the native species have no immunity, modifying the habitat and threatening biodiversity (Bax et al., 2003).

North-western Iberia is a relevant area to study the effects of climate change in the biodiversity and biogeography of intertidal macroalgae because it is a transition zone – representing the rear edge for several cold-temperate species and the leading edge or biogeographical gap for warm-water species. In particular, several cold-water macroalgae typical of higher latitudes find in northern Portugal their last refuge in southern Europe resulting in a regional biodiversity hotspot (Van den Hoek and Donze, 1967). Moreover, historical surveys conducted in the area provide a baseline upon which to quantify and understand ongoing range shifts; work published thus far already indicates changes in the distribution and abundance of intertidal taxa in north-western Iberia (Lima et al., 2006; Wethey et al., 2011; Rubal et al., 2013; Piñeiro-Corbeira et al., 2016; Casado-Amezúa et al., 2019), though not always in the direction expected given recent local patterns of climatic changes (Lima et al., 2007, 2009; Seabra et al., 2015).

From a climatic and oceanographic standpoint, north-western Iberia is remarkable due to the markedly colder summer sea surface temperatures than those found in neighboring regions, a pattern driven by the Canary Upwelling System (Fiuza, 1983; Fraga, 2012). Nevertheless, the region has been warming in recent decades, with evidence suggesting that the buffering effect of upwelling may be slowing down, and forecasts indicating that this region will continue to warm and most likely experience more frequent extreme events in the future (Lima and Wethey, 2012; Seabra et al., 2019). In addition, there is a latitudinal temperature gradient along the Portuguese coast, with warmer waters towards the south. Hence, central Portugal is slightly warmer during winter and less influenced by upwelling during the summer than and northern Portugal and Galicia (north-west Spain, Miranda et al., 2002). Moreover, the transition from central to northern Portugal is marked by a long stretch of sand beaches (∼175 km) interspersed with artificial seawalls and harbors. In turn, while no such physical barrier exists between northern Portugal and Galicia, the north-western coast of Spain has a distinct geomorphology dominated by Rias – large coastal inlets formed by the partial submergence of unglaciated river valleys – compared to a rather rectilinear coastline in northern Portugal, and intense human activity (maritime traffic, ship building, water sports, extensive aquaculture, fishing, among others). Galicia is also a region quite affected by biological invasions (Blanco et al., 2021).

Here we analyze the distribution and abundance of a selected group of macroalgae in north-western Iberia with a fine-scale ad-libitum timed search designed to capture the small-scale variability between sites and robustly identify abundances at or near the distribution limits of selected species. These data were then contrasted against historical surveys in the area (Lima et al., 2007; Pereira et al., 2021a) to assess possible range shifts of warm-water, cold-water and neutral species. Additionally, we also explored if the regions of Galicia, northern Portugal, and central Portugal display distinctive characters regarding macroalgae composition and abundance, and how those characters might have been recently changing.

## 2 Material and Methods

### 2.1 Selection of species and sites

To detail distributional limits of intertidal macroalgae species in northern Portugal and neighboring regions - Galicia and central Portugal - we selected a list of 34 species to be surveyed with high spatial resolution (52 locations over 750 km). We then applied a survey method (ad-libitum timed search with in-situ determination of abundance on a semi-logarithmic scale) appropriate to detect rare species, from our list of selected species (such as cold-water species disappearing from a region or warm-water species recently arriving), in highly variable environments (such as the intertidal).

Species were selected based on previous studies (Veiga et al., 2014; Pereira et al., 2021a) that identified macroalgae with range limits in the area (either rear- or poleward-edge) or that reached neighboring regions and thus can be in the process of migrating into the area facilitated by ongoing environmental changes and/or anthropogenic disturbance (e.g., introduced species, Supplementary Table 1 and Pereira et al. 2022). Importantly, Lima and colleagues have previously described that while warm-water species in this region had been moving northwards, cold-water species had been moving on both directions, with no clear trend (Lima et al., 2007). Hence, to better understand the direction of change and to compare our results with those previous findings, we classified species as having “cold-water”, “warm-water” or “neutral” affinity. The classification was based on how their Species Temperature Index (STI) compared with temperatures in the region - if it was lower, higher, or in-between the range of average temperatures in surveyed sites, respectively. Then, to compare between regions and between historical and current surveys, we computed the average STI of all surveyed species, per site. For details, see Pereira et al. (2022).

Sites surveyed included nearly all wave-exposed rocky shores (38) in our region of interest – northern Portugal (Supplementary Table 2). In addition, four sites in central Portugal and 11 in Galicia (north-west Spain) were used to determine the uniqueness of northern Portugal relative to its neighboring regions, in a total of 52 sites. Twenty-three of the locations had been surveyed in the past (2001-2005) by the same research group (Pereira et al., 2021a, 2021b, 2021c, unpublished data). Furthermore, we sampled 18 additional sites with artificial hard substrate to finely track the ongoing early-stage invasion of the kelp *Undaria pinnatifida* (already reported in 2 locations in Portugal at the beginning of our surveys,Veiga et al., 2014). This subset of sites where all located within large stretches of sandy coast, and some were in proximity of marinas and harbors, *Undaria*’s likely entry points (Table 3 and Pereira et al. 2022).

### 2.3 Data collection

All locations were surveyed during low tides by a two-person team. Each site was surveyed for 60 minutes. Whenever possible due to geographical proximity, the team surveyed two locations during the same low tide. While surveying a site, the team carefully searched for each of the selected species and recorded the abundance based on the SACFOR scale (Southward et al., 1995; Burrows et al., 2008). This scale grades abundances as Superabundant, Abundant, Common, Frequent, Occasional and Rare. Absent species are recorded as Not Found. In the case of locations surveyed more than once during this study, the higher SACFOR abundance was kept in all analyses.

The quadrat sampling methodology is a staple method in ecology, which owing to its inherent objectivity and precision in the assessment of the percentage cover of species, and its easy integration with automated methodologies (e.g., quadrats can be photographed and pictures automatically processed, Bravo et al., 2021) is widely used to quantify the occurrence and abundance of sessile marine species (Boaventura et al., 2002; Bertocci et al., 2012; Franklin et al., 2013). It does not, however, efficiently detect rare species or species inside tide pools or in otherwise hard to reach surfaces (such as rock overhangs, slippery slopes, or areas exposed to the surf during the surveys). In turn, ad-libitum timed search for selected species, quantifying and recording their abundance in a simple, semi-logarithmic scale (i.e., the SACFOR scale, Southward et al., 1995; Burrows et al., 2008) is, at least conceptually, a more appropriate method to pinpoint the distribution limits of species, where it is more likely that populations will occur with low densities or even cryptically in specific microhabitats.

The identification of the selected species was based on Araújo et al. (2009, 2011), Aziza et al. (2008), Bárbara (2009, 2013), Benita et al. (2018), Bunker et al. (2017), Campbell et al. (1994), Chapman & Goudey (1983), Edwards et al. (2012), Faes & Viejo (2003), Molenaar et al. (1996), Poza et al. (2017), Stuart et al. (1999), Vieira et al. (2010). For further details on the species surveyed, locations visited, and specific methodology refer to Pereira et al. (2022).

### 2.4 Data Analysis

Data analysis was divided into (i) the analysis of contemporary data (years 2020 and 2021) from all 52 locations, and (ii) a comparative analysis between present-day data and historical data (recorded between 2001 and 2005) from the 22 locations for which historical data was available (Supplementary Table 2). Data analyses were performed in R (R Core Team, 2020).

#### 2.4.1 Current data (2020 and 2021)

A preliminary analysis was done by creating species distribution and abundance maps using the packages ggplot2, PBSmapping, and ggrepel (Schnute et al., 2004; Wickham et al., 2016; Slowikowski et al., 2018). Coastline information was based on the GSHHG (Global Self-consistent, Hierarchical, High-resolution Geography Database) downloaded from https://www.ngdc.noaa.gov/mgg/shorelines/. Similar maps were created for historical and present-day distribution data of all selected species (Supplementary Table 1). Then, using the packages viridis, ggplot2, reshape2 and hrbrthemes (Wickham, 2012; Wickham et al., 2016; Garnier et al., 2018; Rudis, 2020), a heatmap was created representing the abundance and distribution of all surveyed species in 2020/2021. To facilitate recognition and interpretation of patterns and differences among regions, non-metric multi-dimensional scaling (NMDS) plots were produced using the vegan “metaMDS” function based on Bray-Curtis dissimilarity values, calculated from untransformed data by the function “vegdist”, also from the package vegan (Oksanen et al., 2013). The site “Areia” (ID 27) was not included in this analysis as it was an extreme outlier. Finally, using the “adonis” function from the vegan package (Oksanen et al., 2013), the distinction between regions identified in the NMDS ordination plot was confirmed by means of a PERMANOVA test using the Bray-Curtis dissimilarity values and 999 permutations, followed by pairwise comparisons between each group.

Indicator species for each region (Galicia, northern Portugal, central Portugal) were identified using the function “IndVal.g” from the R package “indicspecies” (De Caceres et al., 2016), which corrects the phi coefficient for the fact that some regions have more sites than others and returns the better matching species patterns, assessing the predictive values of species as indicators of the conditions prevailing within each group of locations. This function also outputs specificity (“A”) and fidelity (“B”) values, the specificity component being the estimate of the probability that the surveyed site belongs to the target site group given the fact that the species has been found, and the fidelity component being the estimate of the probability of finding a given species in sites belonging to the target site group (De Cáceres, 2020).

To test for differences in the averaged STI between the regions of Galicia, northern Portugal, and central Portugal, a Krustal-Wallis test was performed using the “multcomp” package (Hothorn et al., 2016), followed by a pairwise comparison using the Wilcoxon rank sum exact test.

#### 2.4.2 Temporal analysis (historical and present-day data)

NMDS plots were performed to facilitate recognition and interpretation of patterns between historical and present-day data, following the procedure described above. To test for a possible trend towards community homogenization manifested in the reduction of the average Bray-Curtis dissimilarity from historical to present-day data, a permutation test was employed (Petrov, 2021). Indicator species for the historical and current group were also identified using, as before, the function “IndVal.g” from the R package “indicspecies” (De Caceres et al., 2016). A one-way ANOVA test was used to test for differences in the average STI of the selected species, also using R.

## 3 Results

### 3.1 Current data

The heatmap plot in Figure 1 shows an overview of the occurrence and abundance of the 34 species surveyed in the 52 locations in north-western Iberia in 2020/2021. Despite having met the criteria for inclusion in this survey, the species with warm-water affinity *Valonia utricularis, Phyllophora crispa, Hypnea musciformis, Halopithys incurva*, and the cold-water affinity *Phycodrys rubens, Dumontia contorta* and *Delesseria sanguinea* were not found in any of the locations surveyed. Some taxa were commonly found across the whole study area - *Sacchoriza polyschides, Sargassum muticum, Treptacantha baccata, Grateloupia turuturu, Fucus spiralis*, and *Chondrus crispus*. In contrast, some species were only found in a single site, such as *Sargassum flavifolium, Saccharina latissima, Petalonia fascia, Padina pavonica, Halidrys siliquosa, Ascophyllum nodosum* and *Fucus serratus*.

**Figure 1.**
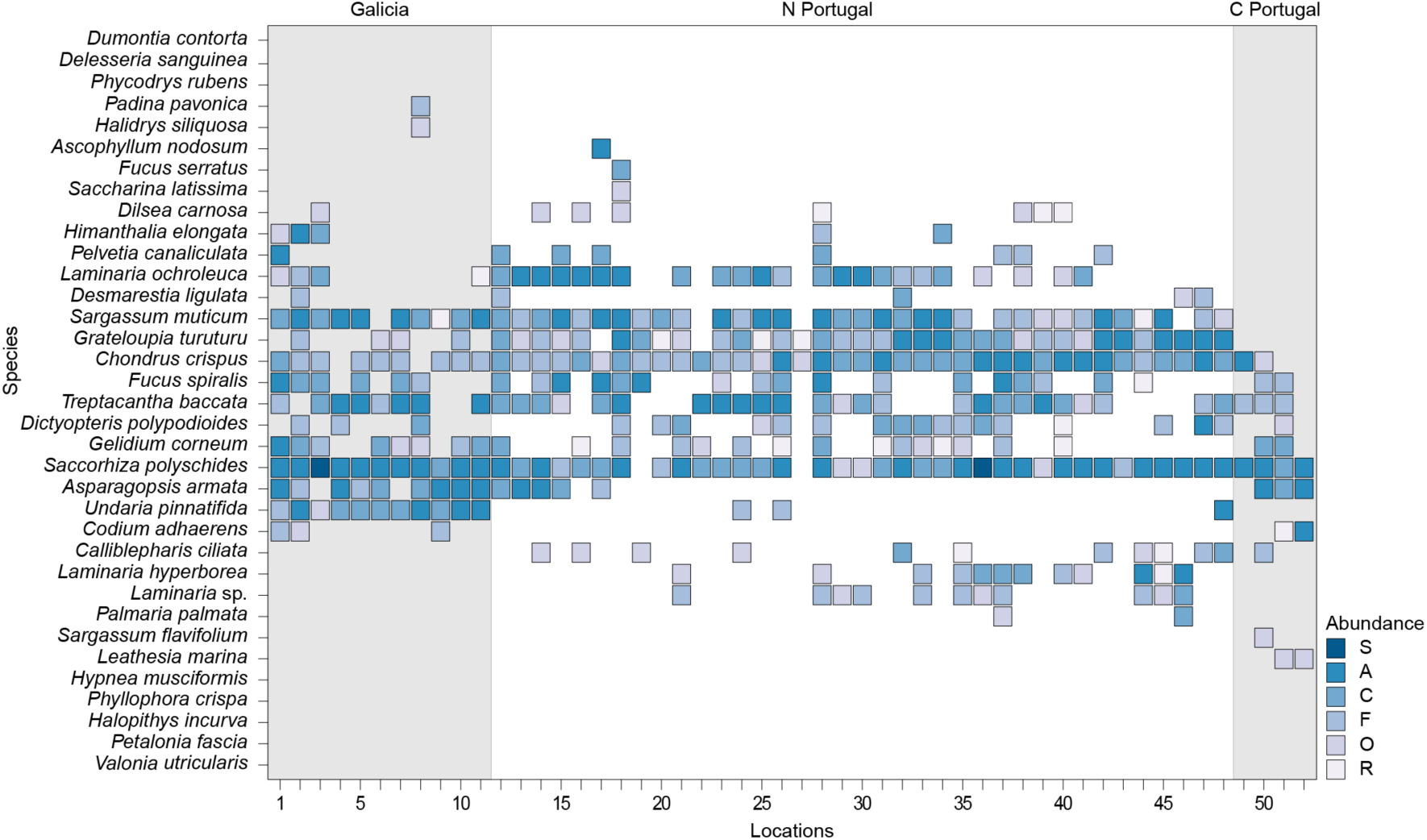
Abundance of selected species (y-axis) in North-western Iberia. In the x-axis, locations are identified by an ID number, from the northernmost (1) to the southernmost site (52). For more details see Supplementary Table 2. Color gradient indicates abundance based on the SACFOR abundance scale.

*Asparagopsis armata* and *Codium adhaerens*, both warm-water species, displayed a prominent gap in their distribution in Northern Portugal, in clear contrast with *Laminaria hyperborea*, a cold-water species, which could only be found in that region.

The NMDS of the contemporary data (Figure 2) revealed a distinct separation in the multidimensional space between sites from the regions of Galicia, northern Portugal, and central Portugal, which displayed significantly different macroalgae communities between every region (PERMANOVA, p = 0.001, Supplementary Tables 3 and 4).

**Figure 2.**
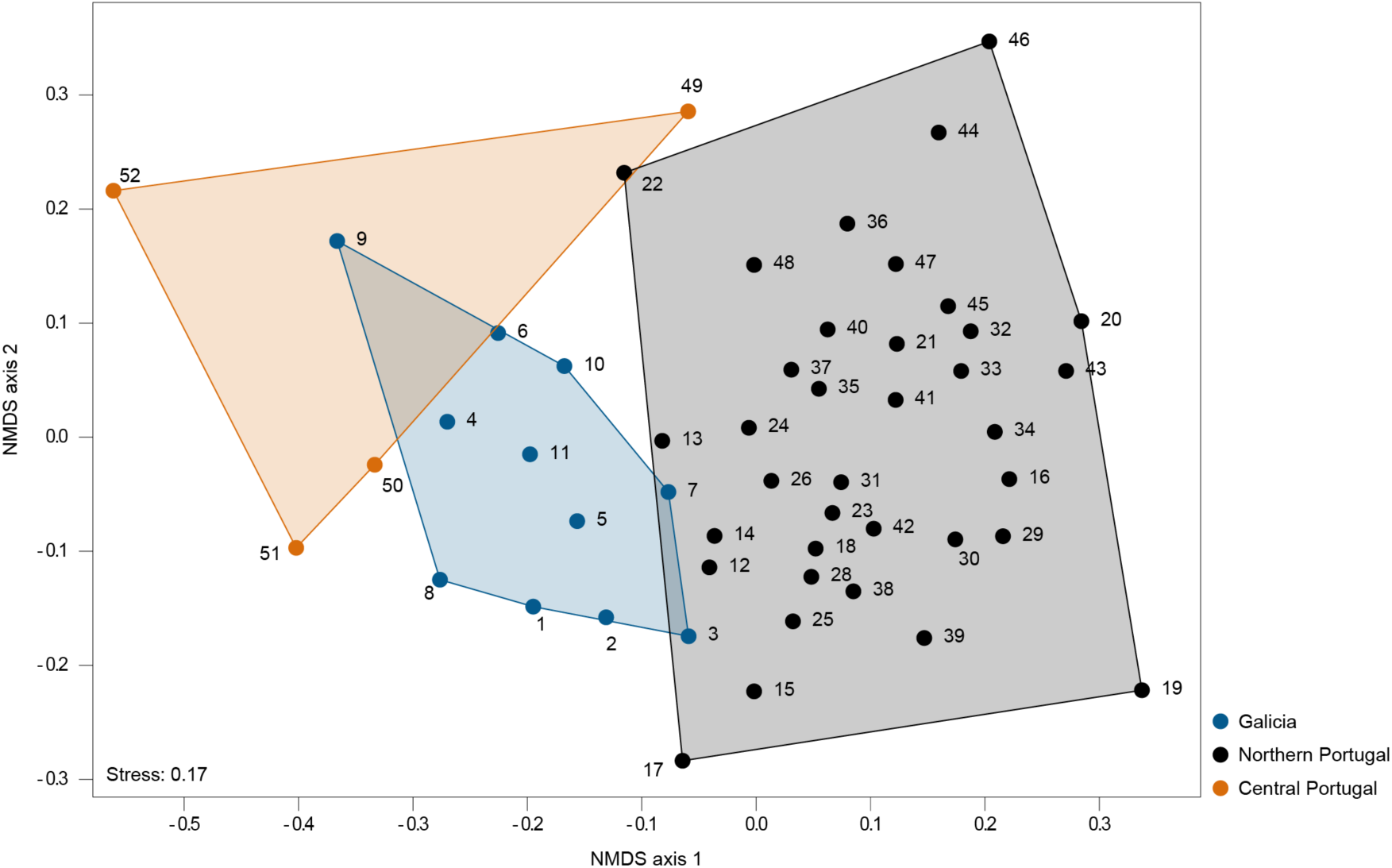
NMDS ordination plot of all sites based on the Bray-Curtis dissimilarity index. Blue symbols and blue polygon correspond to sites in Galicia, black symbols and black polygon correspond to sites in northern Portugal and red symbols and red polygon correspond to sites in central Portugal. Locations are depicted by an ID number, for more details see Supplementary Table 2.

#### 3.1.1 Indicator species

Using a Multilevel Pattern Analysis, we extracted information on the indicator species characteristic of a region or group of regions (Supplementary Table 5). In Galicia, *Undaria pinnatifida* (Supplementary Figure 1B) was the only species significantly characteristic of the region. It showed a high specificity value (A = 0.93), as it occurred almost exclusively within that region, with a few exceptions in northern Portugal. *U. pinnatifida* also displayed a high-fidelity value (B = 1), as it was present in most locations in Galicia that were sampled in this study.

Regarding northern Portugal, the analysis could not identify significant representative species. However, five species, all with cold-water affinity - *Laminaria hyperborea, Palmaria palmata, Ascophyllum nodosum, Fucus serratus*, and *Saccharina latissima* - were flagged as associated with this region because they occurred almost exclusively in northern Portugal (thus receiving high values of specificity - A), although their occurrence was not consistent among all sites (hence, receiving relatively low values of fidelity - B).

The species that significantly separated central Portugal from the other two regions were the cold-water *Leathesia marina* (Supplementary Figure 2B) and the warm-water *Codium adhaerens* (Supplementary Figure 3B), which are specific to this region.

Considering region pairs, both *Sargassum muticum* (Supplementary Figure 4B) and *Grateloupia turuturu* (Supplementary Figure 5B) held high values of specificity (A = 0.91 and 0.81, respectively) in Galicia and northern Portugal, significantly representing both regions. The species significantly associated with Galicia and central Portugal were *Asparagopsis armata* (Supplementary Figure 6B) and *Gelidium corneum* (Supplementary Figure 7B), mostly due to their high values of specificity (A = 0.92 and 0.83 respectively), indicating that they were either rare or non-existent in northern Portugal. Both are warm-water species in the study area. There were no species significantly associated with both northern and central Portugal, and only *Calliblepharis ciliata* (Supplementary Figure 8B) could be said to represent both regions (although non significantly). This indicates that, regarding the surveyed species, these are two distinct regions, a pattern corroborated by the NMDS and multivariate analysis (Figure 2, Supplementary Tables 3 and 4).

#### 3.1.2 Average Species Temperature Index

The average Species Temperature Index (avg STI) was significantly different between regions (Chi square = 8.86, p = 0.011, df = 2, Supplementary Table 6). Galicia and central Portugal exhibited the same average STI (15 °C), significantly higher than that found for northern Portugal (14 °C) (Wilcoxon rank sum exact test, p-value = 0.009; Supplementary Table 7).

### 3.2 Temporal analysis – historical and present-day data

Table 1 summarizes the information on the species that increased, decreased, or maintained their average abundance between historical and present-day surveys. Among those species that increased their distribution, 4 were warm-water species, 6 were cold-water species and 3 were neutral species. In addition, 4 species were invasive. Among the species that decreased their distribution, 3 were warm-water, 2 were neutral species, and 10 were cold-water species. In other words, when compared to past data and considering only species that changed their distribution and abundance within the study area, more cold-water species decreased than increased (10 to 6) while more warm-water species increased than decreased (4 to 3).

**Table 1.**
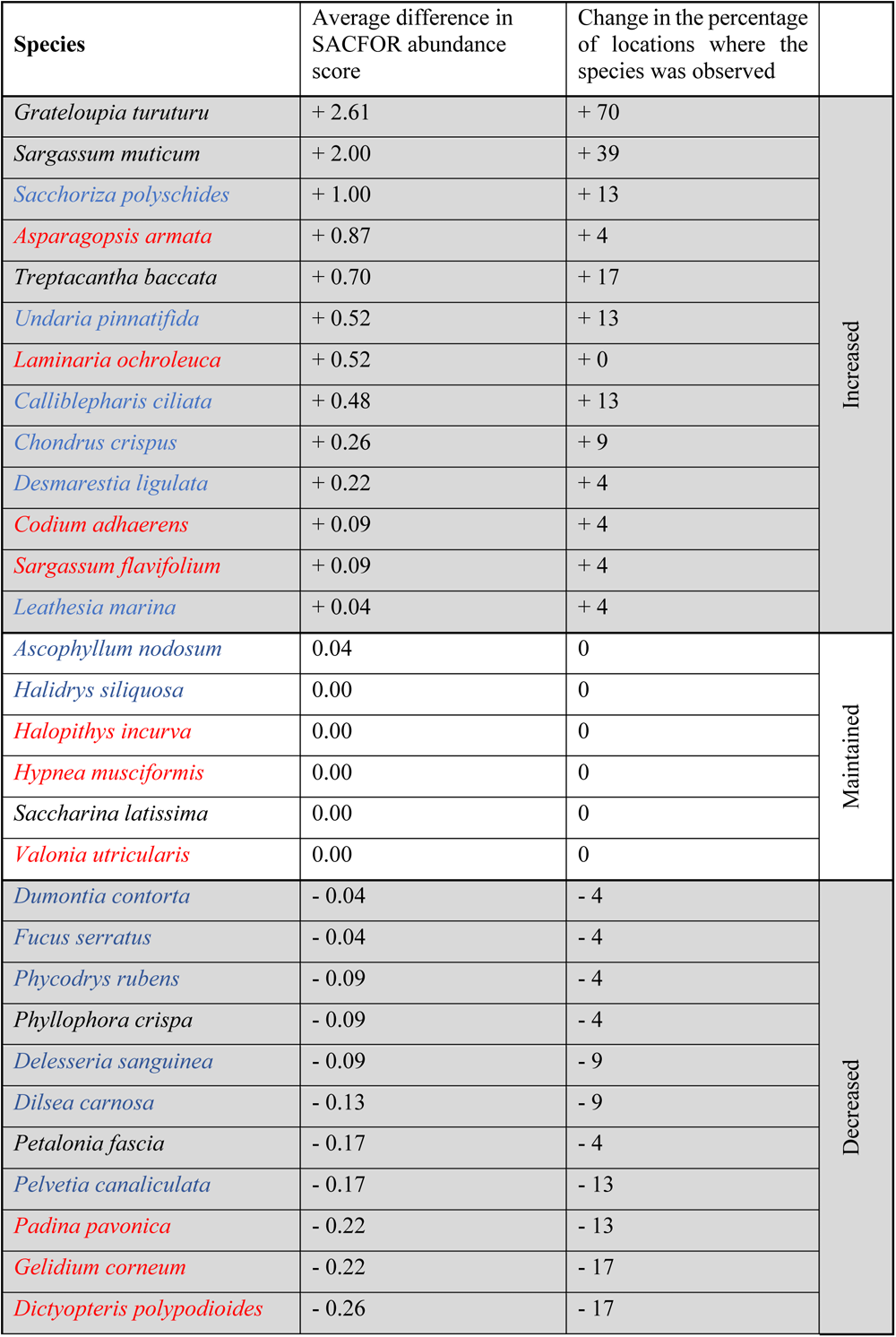
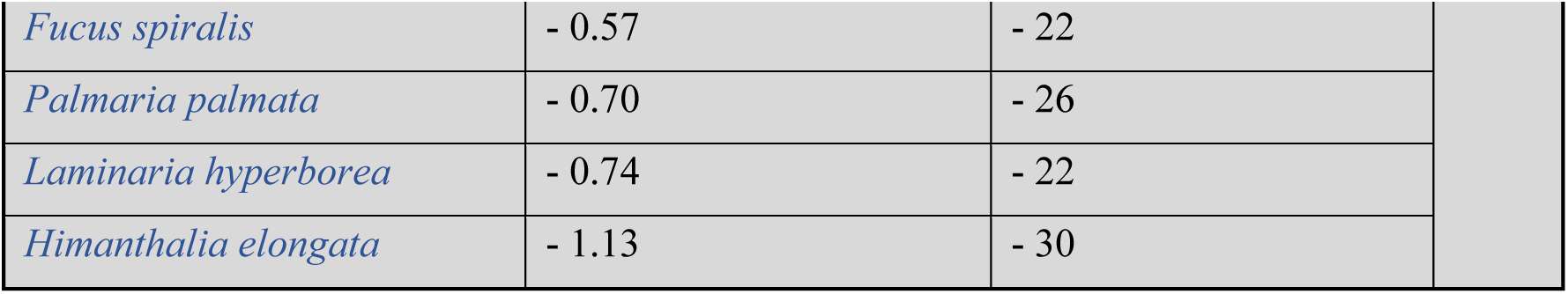
Description of species that increased, decreased, or maintained their average abundance and geographical distribution from historical (2001-2005) to present-day data. Species in blue are cold-water species, while those in red are warm-water species and those in black are neutral species.

| Species | Average difference in SACFOR abundance score | Change in the percentage of locations where the species was observed |  |
| --- | --- | --- | --- |
| *Grateloupia turuturu* | + 2.61 | + 70 | Increased |
| *Sargassum muticum* | + 2.00 | + 39 |  |
| *Sacchoriza polyschides* | + 1.00 | + 13 |  |
| *Asparagopsis armata* | + 0.87 | + 4 |  |
| *Treptacantha baccata* | + 0.70 | + 17 |  |
| *Undaria pinnatifida* | + 0.52 | + 13 |  |
| *Laminaria ochroleuca* | + 0.52 | + 0 |  |
| *Calliblepharis ciliata* | + 0.48 | + 13 |  |
| *Chondrus crispus* | + 0.26 | + 9 |  |
| *Desmarestia ligulata* | + 0.22 | + 4 |  |
| *Codium adhaerens* | + 0.09 | + 4 |  |
| *Sargassum flavifolium* | + 0.09 | + 4 |  |
| *Leathesia marina* | + 0.04 | + 4 |  |
| *Ascophyllum nodosum* | 0.04 | 0 | Maintained |
| *Halidrys siliquosa* | 0.00 | 0 |  |
| *Halopithys incurva* | 0.00 | 0 |  |
| *Hypnea musciformis* | 0.00 | 0 |  |
| *Saccharina latissima* | 0.00 | 0 |  |
| *Valonia utricularis* | 0.00 | 0 |  |
| *Dumontia contorta* | - 0.04 | - 4 | Decreased |
| *Fucus serratus* | - 0.04 | - 4 |  |
| *Phycodrys rubens* | - 0.09 | - 4 |  |
| *Phyllophora crispa* | - 0.09 | - 4 |  |
| *Delesseria sanguinea* | - 0.09 | - 9 |  |
| *Dilsea carnosa* | - 0.13 | - 9 |  |
| *Petalonia fascia* | - 0.17 | - 4 |  |
| *Pelvetia canaliculata* | - 0.17 | - 13 |  |
| *Padina pavonica* | - 0.22 | - 13 |  |
| *Gelidium corneum* | - 0.22 | - 17 |  |
| *Dictyopteris polypodioides* | - 0.26 | - 17 |  |

|  |  |  |
| --- | --- | --- |
| *Fucus spiralis* | - 0.57 | - 22 |
| *Palmaria palmata* | - 0.70 | - 26 |
| *Laminaria hyperborea* | - 0.74 | - 22 |
| *Himanthalia elongata* | - 1.13 | - 30 |

#### 3.2.1 Graphical representation and permutation analysis

The presence and abundance of selected macroalgae species significantly differed between historical (2001-2005) and current (2020-2021) datasets (PERMANOVA, p = 0.001, Supplementary Table 8).

The NMDS presented in Figure 3 shows that the average dispersion of locations around their centroid, based on each site’s species composition, is smaller in current data than in historical data although, probably owing to the high variability within each group, the trend is not statistically significant (t (36.057) = 1.4812, p = 0.1472).

**Figure 3.**
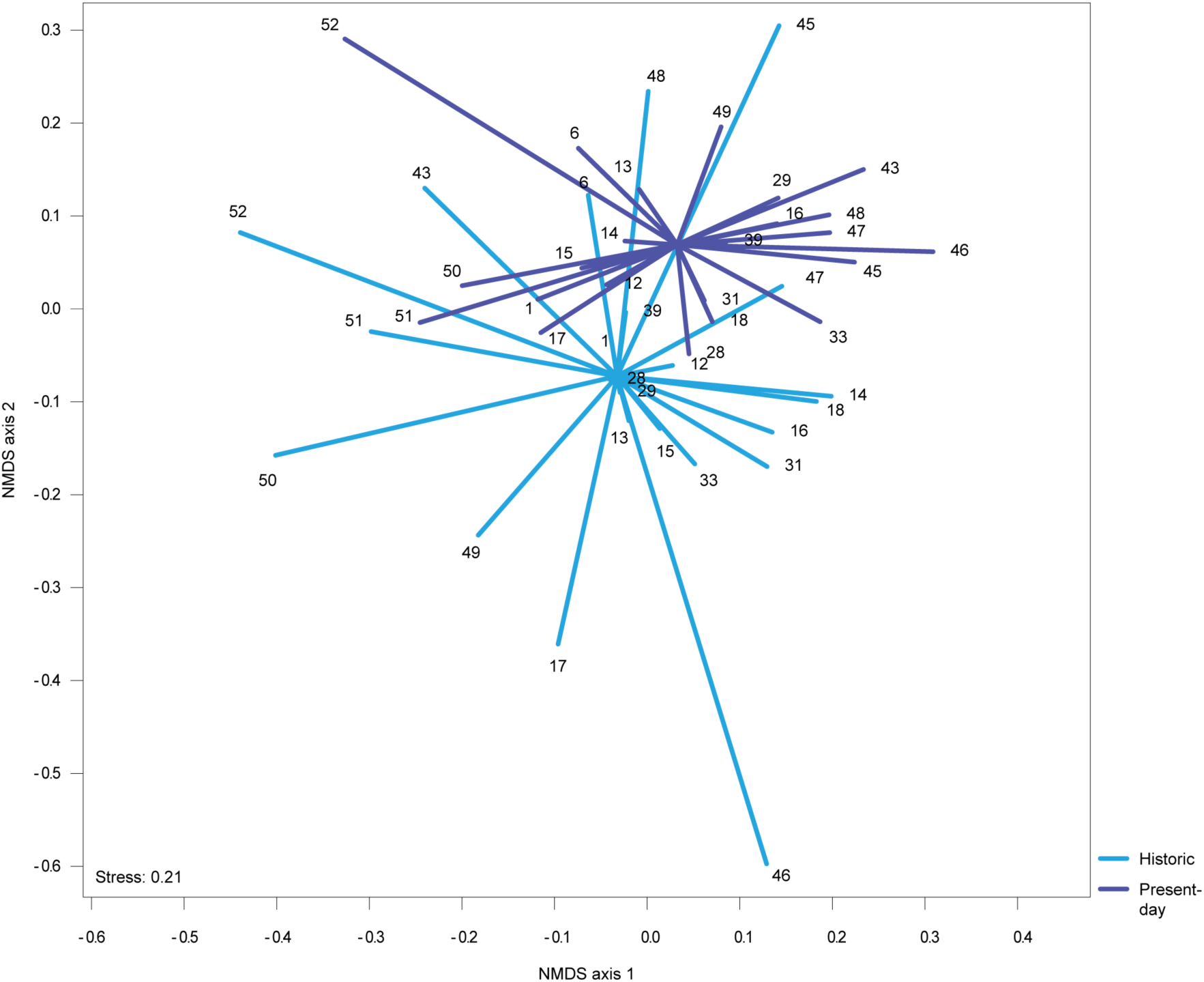
NMDS ordination plot based on the Bray-Curtis dissimilarity index for the list of common sites between present-day (2020-2021) and historical (2001-2005) sampling campaigns. Blue lines connect each historic location to its centroid and the purple lines connect each present-day location to its centroid. Locations are depicted by an ID number, for more details see Supplementary Table 2.

Average fidelity across all sites, computed through indicator species analysis, was higher in 2020-2021 (Average fidelity = 0.383; Standard deviation = 0.335) than in 2001-2005 (Average fidelity = 0.136; Standard deviation = 0.136), which also suggests a tendency towards homogenization of the macroalgae communities, albeit not significant.

#### 3.2.2 Indicator species

*Himanthalia elongata* (Supplementary Figure 9) and *Palmaria palmata* (Supplementary Figure 10) were highlighted as significantly characteristic species in the historical surveys (Supplementary Table 8). Although found at relatively high abundance in numerous locations in Portugal ∼15 years ago, these species have since gone extinct almost everywhere within the study area. Currently, *H. elongata* can only be found at three sites in Galicia and in two sites in Portugal (Praia Central – 34; Mindelo - 28), while *P. palmata* can only be found in a single location in Northern Portugal (Senhor da Pedra - 46).

*Grateloupia turuturu* (Supplementary Figure 5) and *Sargassum muticum* (Supplementary Figure 4) are invasive species that have recently expanded their geographical distribution and increased in abundance in the Portuguese coast, resulting in their significant contribution to the 2020/2021 group (Supplementary Table 8).

#### 3.2.3 Average Species Temperature Index

The analysis revealed that most locations have increased their average STI from 2001-2005 to 2020-2021 (i.e., locations above the y = x line in Figure 4). Every location either maintained or increased their average STI except for the sites Senhor da Pedra (ID-46) and São Martinho do Porto (ID-50). Still, this trend was not significant overall (F (1,44) = 0.947, p = 0.34).

**Figure 4.**
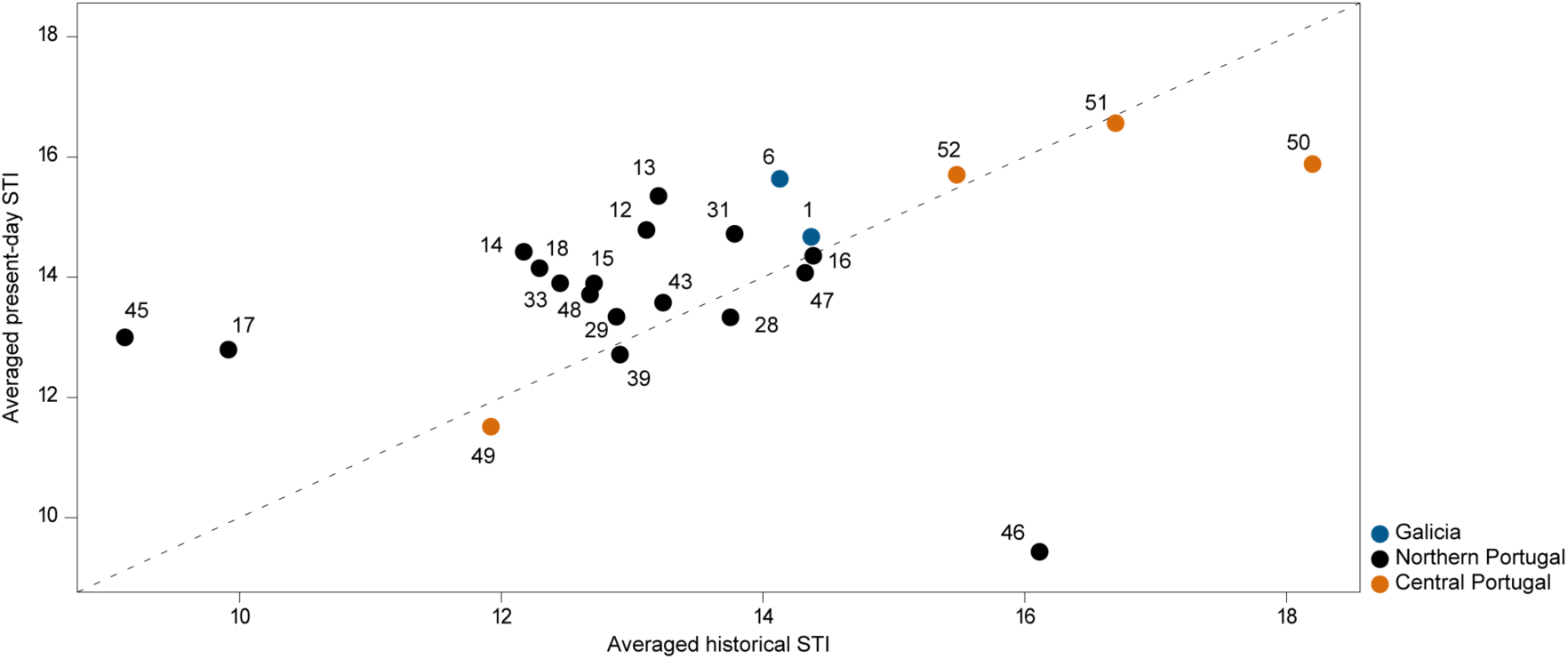
Comparison between averaged Species Temperature Index of sites in the present (averaged present-day STI) with community temperature index of sites in the past (averaged historical STI). Red circles represent sites in central Portugal, black circles represent sites in northern Portugal and blue circles represent sites in Galicia. The line y = x shows the theoretical position of points if there were no changes in average STI. Locations are depicted by an ID number, for more details see Supplementary Table 2.

## 4 Discussion

### 4.1 Fine-scale sampling

We used a fine-scale survey strategy that allowed us to obtain distribution and abundance data at an average resolution of 3.4 ± 5.9 km within the region of highest interest (northern Portugal), a value higher than what is typically provided in similar studies (Boaventura et al., 2002; Araújo et al., 2009). Such high-resolution distribution data proved instrumental, particularly in circumstances where certain species were found to have disappeared from historical sites only to be recorded in other locations just a few hundreds of meters away (e.g, *Pelvetia canaliculata*, which disappeared from Cabo do Mundo (ID-39) but was found 2.2 kms away at site Memória (ID-38)). This means that the most common long-term monitoring strategy, which is to target all the effort towards revisiting the same sites repeatedly (Mieszkowska et al., 2021), may yield oversimplified datasets, and be insufficient to accurately pinpoint the distribution limits of certain species or to detect and therefore track fast changes. Moreover, previous studies have already demonstrated that species distribution models based on fine-scale surveys result in different patterns from those fed with data obtained from coarser surveys (Trivedi et al., 2008; Gastón and García-Viñas, 2010), yielding qualitatively distinct predictions with important consequences for the conservation and management of biodiversity (Franklin et al., 2013).

By adopting the ad-libitum timed search for selected species and recording their abundance in a simple, semi-logarithmic scale (i. e., the SACFOR scale, Southward et al., 1995; Burrows et al., 2008), we successfully mitigate the drawbacks associated with the commonly-used quadrat sampling method. In particular, while the combined ad-libitum timed search with SACFOR recording can be regarded as a less objective or accurate method which often requires expert knowledge (and thus is harder to train for, Mieszkowska et al., 2006), our study confirms that it can efficiently detect rare species or species in hard-to-reach microhabitats, unlike the quadrat method. Furthermore, our data strongly suggests that the use of intensive, fine-scale surveys was essential to accurately assess the distribution of the studied species, which in turn allowed us to formally test hypotheses on effects of warming on species abundances at or near their range edges. This clearly highlights the importance of taking into consideration the overall goals of any assessment of intertidal biodiversity when choosing the spatial coverage required and the sampling method to be used.

### 4.2 Regional variation in the north-western Iberian Peninsula

Considering macroalgae composition and abundance, our results show that the regions of Galicia, northern Portugal and central Portugal are all significantly different from each other. The distinction between central Portugal and other regions was expected, since it is geographically isolated by a long stretch of sand and is less affected than the other regions by the seasonal upwelling of cold and nutrient-rich waters (Fiuza, 1983; Lima et al., 2007; Fraga, 2012; Seabra et al., 2015). Previous studies on the distribution of intertidal species along the Portuguese coast had also classified northern and central Portugal as distinct biogeographic regions (Van den Hoek and Donze, 1967; Boaventura et al., 2002; Lima et al., 2007).

Remarkably, this study also suggests that Galicia and northern Portugal are significantly different in their seaweed composition and abundance despite being geographically very close (they are part of the same continuum of rocky coast) and having (at least apparently, and based on remote sensing data) similar sea surface temperature profiles (Banzon et al., 2016). The underlying reason for these differences is not evident as the two regions have rarely been surveyed and compared in a standardized way until now. One hypothesis is that macroalgae communities in Galicia and northern Portugal are distinct due to small-scale temperature variability which is not easily detected from remotely sensed data due to its insufficient resolution (Meneghesso et al., 2020). In reality, it would be surprising if the contrasting topography and geomorphology contexts associated with each of the regions would not create dissimilar environmental conditions, not only in relation to water and atmospheric temperatures, but also regarding water salinity, wave exposure, tidal regime, current speed, nutrient availability, or magnitude of upwelling, all of which have the potential to influence local biodiversity. Specifically, the coast of Galicia is characterized by the occurrence of Rias (large coastal inlets formed by the partial submergence of unglaciated river valleys), while the coast of Northern Portugal is linear, with a north to south orientation, and absent of sheltered sites.

Another hypothesis is that the differences detected could be related with the different anthropogenic pressures exerted in these regions. The aquaculture production of seaweed and bivalves (Minchin and Nunn, 2014; James and Shears, 2016) and the heavy traffic of leisure and fishing vessels in the Rias might represent vectors for biological invasions (Blanco et al., 2021). Several invasive species have higher local abundances and are distributed more widespread in Galicia than in northern Portugal (e.g., *Undaria pinnatifida* -Supplementary Figure 1 and *Asparagopsis armata* – Supplementary Figure 6).

The species *Undaria pinnatifida* and *Himanthalia elongata* are indicator species in Galicia, while occurring in just a few sites in Portugal. They are both cold-water species with their southern distributional limit in the north-western Iberian Peninsula. These seaweeds have, however, different geographical origins. The invasive *U. pinnatifida* is common in Galicia and is currently spreading along the northern Portuguese coast (this study; Báez et al. 2010; Veiga et al. 2014). Contrastingly, the native *H. elongata*, once abundant in the study area, has been retreating from its historical meridional distribution limit in central Portugal (Ardré, 1971) and decreasing in abundance and distribution both in Galicia and northern Portugal (this study; Barrientos et al. 2020; Piñeiro-Corbeira et al. 2016; Ramos et al. 2020). Both species are commercially harvested and cultivated in Galicia which might explain the maintenance of relatively larger populations in Galicia than in northern Portugal, resulting from spill-over from aquaculture production (Lagos and Cremades, 2004; Minchin and Nunn, 2014).

Interestingly, Galicia and central Portugal have a higher average species temperature index than northern Portugal. Additionally, all indicator species that significantly represent both Galicia and central Portugal are considered warm-water species in Northern Portugal (*Asparagopsis armata* and *Gelidium corneum*), thus supporting the hypothesis that temperature may play an important role in the patterns here described. Our results suggest that Northern Portugal is an important thermal refugia for several cold-water species (e.g. *Laminaria hyperborea, Ascophyllum nodosum* or *Fucus serratus*) which, considering the surveyed locations, were only found there. Nevertheless, for some species such as *Himanthalia elongata* or *Halidrys siliquosa*, the potential thermal refugia of this region seems to be eroding. It will soon be possible to test this hypothesis through the analysis of high-resolution intertidal temperature profiles currently being collected by autonomous temperature loggers (Lima and Wethey, 2012) at the same locations where the biodiversity surveys were carried out.

There is high variability in species composition and abundance in northern Portugal, as revealed by the lack of significant representative species (Supplementary Table 5) and large dispersion among sites within the region (Figure 2). Notwithstanding the fact that the sampling effort in this region was much higher, the within-region biodiversity variability suggests that even a seemingly linear and homogeneous coastline holds significant high-frequency spatial variability. One explanation for this pattern is that biodiversity variability is simply mirroring the micro-climate variability created by subtle, yet biologically important differences in solar radiation exposure, salinity, topography, wave exposure and hydrodynamics (Seabra et al., 2011; Lima et al., 2016). Combinations of these different environmental conditions create a complex mosaic of micro-habitats, even among neighboring locations, which might explain the high biodiversity variability (Helmuth et al., 2006; Potter et al., 2013). Hence, the northern Portuguese coast is a good case study to explore how the effects of small-scale environmental variability interact with large-scale patterns, namely global changes in sea temperature.

In conclusion, our analyses indicate that the regions of Galicia and central Portugal, although geographically separated, share a regionally warm-water species pool. In turn, Northern Portugal is, at least from the standpoint of the average Species Temperature Index and from the distribution of cold-water species, the coldest region, highlighting its potential importance as a climate change refugia that may be sustaining the equatorial range edge of several macroalgae. This calls for protection of the rocky intertidal in this region since climate change refugia are increasingly considered in conservation priorities (Morelli et al., 2020).

### 4.3 Temporal variation between present-day and historical data

Macroalgae occurrence and abundance have changed considerably in the 15 years that mediate our historical (2000-2005) and contemporary surveys (2020-2021). Currently, macroalgae communities seem to be more homogeneous, although the effect is small and not significant. This homogenization, also evident in subtidal macroalgae communities along the coast of Portugal (de Azevedo, 2019), could lead to impacts in ecosystem functioning, productivity and ecosystem services (Clavel et al., 2011).

In addition, we observed a net decrease in abundance and distribution of cold-water species and a net increase of warm-water and neutral species. This contrasts with results from Lima et al. (2007), that observed a clear trend of northward migration of warm-water species and mixed responses of cold-water species (some were retreating northwards while others were advancing southwards). Given that this work comprises two datapoints in time separated by 15 years, and the reduced geographical area surveyed when compared with Lima et al. (2007), we cannot confirm that a shift in the overall direction of change in communities is occurring. Still, our results are concerning especially given the accentuated decrease in some cold-water (foundation) species and the parallel increase of some warm-water (invasive) macroalgae, both with the potential to disproportionately impact the entire ecosystem.

Several macroalgae for which we report decreases in presence and abundance are canopy-forming brown algae that play an important role in coastal biodiversity as foundation species (sensu Dayton, 1972), providing food, shelter, and habitat to other species of flora and fauna, as well as protection from desiccation, temperature extremes, and wave action (Bulleri et al., 2006; Smale et al., 2013; Seitz et al., 2014). Similar declines have been reported in other regions of the Atlantic, notably the British Isles (Yesson et al., 2015). This is the case for *Himanthalia elongata* (Creed, 1995) which, among all the species here surveyed, and in agreement with previous studies (Piñeiro-Corbeira et al., 2016; Casado-Amezúa et al., 2019; Barrientos et al., 2020), showed the greatest populational and geographic declines in Galicia and Portugal (a 30% decrease in the number of sites and an average 1.13 reduction in SACFOR score). Currently, it can only be found in two locations south of the NW corner of the Iberian Peninsula - one healthy population in Praia Central (ID - 34) and a very small population in Mindelo (ID - 28) with just a few scattered individuals (Supplementary Figure 9). Similarly, a noticeable decrease of the foundation species *Laminaria hyperborea* was also evident (a 22% decrease in the number of sites and an average 0.74 reduction in SACFOR score, Supplementary Figure 11). This kelp forms marine forests and sustains a diverse community along the north-east Atlantic coast (Christie et al., 2003; Norderhaug et al., 2021). In parallel, the warm-water counterpart *L. ochroleuca* increased slightly in abundance, though not enough to replace *L. hyperborea*. It is important to note that even if successful in replacing *L. hyperborea, L. ochroleuca* supports a significantly less diverse community and has lower biomass (Teagle and Smale, 2018).

In addition, the decrease of *Laminaria hyperborea* could have contributed to the decrease of *Palmaria palmata* (Supplementary Figure 10), as this red algae occurs often as an epiphyte of *L. hyperborea* towards the rear edge of its distribution (Whittick, 1983). If confirmed, this would be a prime example of how changes in the distribution of one species impact another, having cascading effects in the community. The decrease in these canopy-forming algae could lead to negative ecological and economic outcomes since they simultaneously are at the basis of the food-web and are cultivated and harvested with applications in several industries (Plaza et al., 2008). Moreover, populations at the rear-edge are often relict populations, holding unique genetic diversity that might be key for a species’ persistence under current global change trends (Provan and Maggs, 2012).

Contrastingly, some species highlighted in this study as indicator species have recently increased their distributional range and abundance in the region. The invasive warm-water macroalgae *Grateloupia turuturu* (Supplementary Figure 5) and the invasive, neutral species *Sargassum muticum* (Supplementary Figure 4) have both greatly increased their distribution in the last 15 years. *S. muticum* exhibits extremely high growth rates during spring and is a strong competitor - limiting the distribution and replacing native species, which once established can have a drastic impact in ecosystem flora and fauna composition (Critchley et al., 1986; Boudouresque et al., 1995; Stæhr et al., 2000; Sanchez and Fernandez, 2005). Additionally, the kelp *Undaria pinnatifida*, an invasive cold-water species which was never recorded during the surveys of 2000-2005 and since then had only been recorded in Portugal in Buarcos (ID - 48) and within a marina in Viana do Castelo (Báez et al., 2010; Veiga et al., 2014), was now observed to be spreading rather quickly. It is currently present in four more sites, two with artificial substrate (a breakwater in Cabedelo and the northern breakwater at Barra), and two with natural rocky substrate (24 – Carvalhido and 26 – Forte São João). This species has a year-round presence in non-native areas, where it competes with native species with seasonal reproductive, growth and senescence stages, resulting in a successful establishment and spread in cold to temperate systems (Epstein and Smale, 2017). It is considered one of the top-100 worst invaders (Lowe et al., 2000), causing severe ecological damages in some regions, however with rather neutral and even benign effects in others (South et al., 2017), making it very difficult to predict ecosystem-wide effects of its current spreading in north-western Iberia.

The dramatic changes in the distribution of some invasive species are cause for concern and call for a close and continuous monitoring of these macroalgae in north-western Iberia. Considering that their increase in abundance led to significant ecological disturbance in other regions (Stæhr et al., 2000; Casas et al., 2004; Sanchez and Fernandez, 2005; Williams and Smith, 2007; Silva et al., 2021), it is likely that they will too significantly and negatively impact ecosystem functioning in north-western Iberia (Mineur et al., 2015). Moreover, invasive species, such as *G. turuturu* in northern Portugal (Araújo et al., 2011) could also be taking the role of passenger of ecological change rather than driver, benefiting from the loss of algae canopies and relying on the disruption of native assemblages to spread (Mulas and Bertocci, 2016), in which case, these species can be important indicators of ecosystem health.

The average Species Temperature Index (STI) has increased in several sites, although not significantly. A different study including more macroalgae species has shown that Community Temperature Index (CTI) is increasing as sea surface temperature increases, though the CTI of animals is responding more to changes in temperature than that of macroalgae (Burrows et al., 2020), which might explain the marginal increase in the average STI in our study. However, these change in average STI are congruent with the results mentioned previously, and match findings from other studies, where the disappearance of cold-water species due to warming has caused an increase in CTI (Mclean et al., 2021).

In conclusion, the contrasts between historical and present-day biodiversity and biogeography patterns suggest that a homogenization of the macroalgae communities may be underway across the north-western Iberia. We found that the average community dissimilarity seems to have been decreasing over the last two decades, both within and among regions. Some cold-water species are becoming rarer while warm-water counterparts are found over larger spans of coast. Another factor of homogenization is the increasing prevalence of some invasive species, which already dominate some microhabitats and/or sites. Hence, a higher effort in conservation and restoration is vital to protect the health of these coastal ecosystems and the communities that depend on them (García Molinos et al., 2016). Although this work did not formally explore the role ongoing changes in temperature may be playing in these alterations, both the net decline of cold-water species and the net increase in warm-water species, as well as the higher prevalence of invasive species is congruent with the effects expected from an increase in temperature driven by climate change. These effects could soon be exacerbated by any decrease in the magnitude of the upwelling in the region, which mitigates further warming (Seabra et al., 2019; Sousa et al., 2020). Further studies that explicitly analyze the joint change in biodiversity and temperature are urgently needed to test some of the hypotheses here discussed.

## Author Contributions

Cátia Monteiro: Fieldwork, Data Curation, Data Analyses, Writing - Original Draft and Review and Editing, Supervision.

Joana Pereira: Fieldwork, Data Curation, Data Analyses, Writing – Original Draft and Review and Editing.

Rui Seabra: Data Curation, Writing - Review and Editing, Funding acquisition.

Fernando P. Lima: Conceptualization, Writing - Review and Editing, Funding acquisition, Supervision, Project administration.

## Acknowledgments

We thank Francisco Arenas Parra for his support in taxonomical identification and for revising the manuscript. We thank José Carneiro for his support during fieldwork in Galiza.

## Data Availability Statement

The datasets analyzed for this study can be found in the GBIF - Global Biodiversity Information Facility [http://ipt.gbif.pt/ipt/resource?r=ibpc; http://ipt.gbif.pt/ipt/resource?r=2021_iberianpeninsula; http://ipt.gbif.pt/ipt/resource?r=herbarium].

## References

Antão, L. H., Bates, A. E., Blowes, S. A., Waldock, C., Supp, S. R., Magurran, A. E., et al. (2020). Temperature-related biodiversity change across temperate marine and terrestrial systems. Nat. Ecol. Evol. 4, 927–933. doi:10.1038/s41559-020-1185-7.

Araújo, R., Barbara, I., Tibaldo, M., Berecibar, E., Tapia, P. D., Pereira, R., et al. (2009). Checklist of benthic marine algae and cyanobacteria of northern Portugal. Bot. Mar. 52, 24–46.

Araújo, R., Violante, J., Pereira, R., Abreu, H., Arenas, F., and Sousa-Pinto, I. (2011). Distribution and population dynamics of the introduced seaweed Grateloupia turuturu (Halymeniaceae, Rhodophyta) along the Portuguese coast. Phycologia 50, 392–402. doi:10.2216/10-65.1.

Ardré, F. (1971). Contribution à l’étude des algues marines du Portugal II. Ecol. Chorologie., 359–574.

Aziza, M., Givernaud, T., Chikhaoui-khay, M., and Bennasser, L. (2008). Seasonal variation of the growth, chemical composition and carrageenan extracted from Hypnea musciformis (Wulfen) Lamouroux harvested along the Atlantic coast of Morocco. Sci. Res. Essays 3.

Báez, J. C., Olivero, J., Peteiro, C., Ferri-Yáñez, F., Garcia-Soto, C., and Real, R. (2010). Macro-environmental modelling of the current distribution of Undaria pinnatifida (Laminariales, Ochrophyta) in northern Iberia. Biol. Invasions 12, 2131–2139. doi:10.1007/s10530-009-9614-1.

Banzon, V., Smith, T. M., Chin, T. M., Liu, C., and Hankins, W. (2016). A long-term record of blended satellite and in situ sea-surface temperature for climate monitoring, modeling and environmental studies. Earth Syst. Sci. Data 8, 165–176.

Bárbara, I. (2009). Especies invasoras en Galicia e Introducción a Súa Problemática: Medio Mariño, Flora. Universidad de A Coruña.

Bárbara, I. (2013). Algas marinas y salores de Galicia y norte de España: Parte 2. Laboratorio de Algas Marinas, Facultad de Ciencias, Universidad de A Coruña.

Barrientos, S., Barreiro, R., Cremades, J., and Piñeiro-Corbeira, C. (2020). Setting the basis for a long-term monitoring network of intertidal seaweed assemblages in northwest Spain. Mar. Environ. Res. 160, 105039. doi:https://doi.org/10.1016/j.marenvres.2020.105039.

Bax, N., Williamson, A., Aguero, M., Gonzalez, E., and Geeves, W. (2003). Marine invasive alien species: a threat to global biodiversity. Mar. Policy 27, 313–323. doi:https://doi.org/10.1016/S0308-597X(03)00041-1.

Benita, M., Dubinsky, Z., and Iluz, D. (2018). Padina pavonica : Morphology and Calcification Functions and Mechanism. Am. J. Plant Sci. 09, 1156–1168. doi:10.4236/ajps.2018.96087.

Bertocci, I., Dominguez, R., Freitas, C., and Sousa-Pinto, I. (2012). Patterns of variation of intertidal species of commercial interest in the Parque Litoral Norte (north Portugal) MPA: Comparison with three reference shores. Mar. Environ. Res. 77, 60–70. doi:https://doi.org/10.1016/j.marenvres.2012.02.003.

Blanco, A., Larrinaga, A. R., Neto, J. M., Troncoso, J., Méndez, G., Domínguez-Lapido, P., et al. (2021). Spotting intruders: Species distribution models for managing invasive intertidal macroalgae. J. Environ. Manage. 281, 111861. doi:https://doi.org/10.1016/j.jenvman.2020.111861.

Boaventura, D., Ré, P., Cancela da Fonseca, L., and Hawkins, S. J. (2002). Intertidal Rocky Shore Communities of the Continental Portuguese Coast: Analysis of Distribution Patterns. Mar. Ecol. 23, 69–90. doi:10.1046/j.1439-0485.2002.02758.x.

Boudouresque, C. F., Meinesz, A., Ribera, M. A., and Ballesteros, E. (1995). Spread of the green alga Caulerpa taxifolia (Caulerpales, Chlorophyta) in the Mediterranean: possible consequences of a major ecological event. Sci. Mar., 21– 29.

Bravo, G., Moity, N., Londoño-Cruz, E., Muller-Karger, F., Bigatti, G., Klein, E., et al. (2021). Robots Versus Humans: Automated Annotation Accurately Quantifies Essential Ocean Variables of Rocky Intertidal Functional Groups and Habitat State. Front. Mar. Sci. 8, 1366.

Bulleri, F., Eriksson, B. K., Queirós, A., Airoldi, L., Arenas, F., Arvanitidis, C., et al. (2018). Harnessing positive species interactions as a tool against climate-driven loss of coastal biodiversity. PLOS Biol. 16, e2006852. Available at: https://doi.org/10.1371/journal.pbio.2006852.

Bunker, F. S. D., Maggs, C. A., Brodie, J. A., and Bunker, A. R. (2017). Seaweeds of Britain and Ireland., ed. S. Edition Plymouth, Uk: Wild Nature Press.

Burrows, M., Harvey, R., and Robb, L. (2008). Wave exposure indices from digital coastlines and the prediction of rocky shore community structure. Mar. Ecol. Prog. Ser. 353, 1–12. doi:10.3354/meps07284.

Burrows, M. T., Hawkins, S. J., Moore, J. J., Adams, L., Sugden, H., Firth, L., et al. (2020). Global-scale species distributions predict temperature-related changes in species composition of rocky shore communities in Britain. Glob. Chang. Biol. 26, 2093–2105.

Campbell, A., Selvagens, F.-F. paraa P. dos A., Múrias, A., Santos, P. T., and Soares, M. (1994). Fauna e Flora do Litoral de Portugal e Europa. FAPAS.

Casado-Amezúa, P., Araújo, R., Bárbara, I., Bermejo, R., Borja, Á., Díez, I., et al. (2019). Distributional shifts of canopy-forming seaweeds from the Atlantic coast of Southern Europe. Biodivers. Conserv. doi:10.1007/s10531-019-01716-9.

Casas, G., Scrosati, R., and Luz Piriz, M. (2004). The Invasive Kelp Undaria Pinnatifida (Phaeophyceae, Laminariales) Reduces Native Seaweed Diversity in Nuevo Gulf (Patagonia, Argentina). Biol. Invasions 6, 411–416. doi:10.1023/B:BINV.0000041555.29305.41.

Chapman, A. R. O., and Goudey, C. L. (1983). Demographic study of the macrothallus of Leathesia difformis (Phaeophyta) in Nova Scotia. Can. J. Bot. 61, 319–323. doi:10.1139/b83-035.

Christie, H., Jorgensen, N. M., Norderhaug, K. M., and Waage-Nielsen, E. (2003). Species distribution and habitat exploitation of fauna associated with kelp (Laminaria hyperborea) along the Norwegian coast. J. Mar. Biol. Assoc. UK 83, 687–699.

Clavel, J., Julliard, R., and Devictor, V. (2011). Worldwide decline of specialist species: toward a global functional homogenization? Front. Ecol. Environ. 9, 222–228.

Creed, J. C. (1995). Spatial dynamics of a Himanthalia elongata (Fucales, Phaeophyta) population. J. Phycol. 31, 851–859. doi:https://doi.org/10.1111/j.0022-3646.1995.00851.x.

Critchley, A. T., Farnham, W. F., and Morrell, S. L. (1986). An account of the attempted control of an introduced marine alga, Sargassum muticum, in southern England. Biol. Conserv. 35, 313–332.

Dayton, P. K. (1972). Toward an understanding of community resilience and the potential effects of enrichments to the benthos at McMurdo Sound, Antarctica. in Proceedings of the colloquium on conservation problems in Antarctica (Allen Press Lawrence, Kansas, USA), 81–96.

de Azevedo, J. (2019). What has changed in the macroalgal communities of the Portuguese coast over a 6-year period? Master.

De Cáceres, M. (2020). How to use the indicspecies package (ver. 1.7.8).

De Caceres, M., Jansen, F., and De Caceres, M. M. (2016). Package ‘indicspecies.’ indicators 8, 1.

Edwards, M., Hanniffy, D., Heesch, S., Harnández-kantún, J., Moniz, M., Quéguineus, B., et al. (2012). Macroalgae Fact-sheets., ed. A. Soler-Vila Moniz, M. Ryan Institute.

Epstein, G., and Smale, D. A. (2017). Undaria pinnatifida: A case study to highlight challenges in marine invasion ecology and management. Ecol. Evol. 7, 8624–8642. doi:https://doi.org/10.1002/ece3.3430.

Faes, V. A., and Viejo, R. M. (2003). Structure and dynamics of a population of Palmaria palmata (Rhodophyta) in northern Spain. J. Phycol. 39, 1038–1049. doi:https://doi.org/10.1111/j.0022-3646.2003.02-142.x.

Fiuza, A. F. G. (1983). Upwelling Patterns off Portugal. NATO Conf. Ser. 4 Mar. Sci. 10 A, 85–98. doi:10.1007/978-1-4615-6651-9_5.

Fraga, F. (2012). “Upwelling off the Galician Coast, Northwest Spain,” in Coastal Upwelling (American Geophysical Union (AGU)), 176–182. doi:10.1029/CO001P0176.

Franklin, J., Davis, F. W., Ikegami, M., Syphard, A. D., Flint, L. E., Flint, A. L., et al. (2013). Modeling plant species distributions under future climates: how fine scale do climate projections need to be? Glob. Chang. Biol. 19, 473–483.

García Molinos, J., Halpern, B. S., Schoeman, D. S., Brown, C. J., Kiessling, W., Moore, P. J., et al. (2016). Climate velocity and the future global redistribution of marine biodiversity. Nat. Clim. Chang. 6, 83–88. doi:10.1038/nclimate2769.

Garnier, S., Ross, N., Rudis, B., Sciaini, M., and Scherer, C. (2018). viridis: Default Color Maps from ‘matplotlib.’ R Packag. version 0.5 1, 2018.

Gastón, A., and García-Viñas, J. I. (2010). Updating coarse-scale species distribution models using small fine-scale samples. Ecol. Modell. 221, 2576–2581. doi:https://doi.org/10.1016/j.ecolmodel.2010.07.016.

Hampe, A., and Petit, R. J. (2005). Conserving biodiversity under climate change: the rear edge matters. Ecol. Lett. 8, 461–467. doi:10.1111/j.1461-0248.2005.00739.x.

Harley, C. D. G., Anderson, K. M., Demes, K. W., Jorve, J. P., Kordas, R. L., Coyle, T. A., et al. (2012). Effects of climate change on global seaweed communities. J. Phycol. 48, 1064–1078. doi:10.1111/j.1529-8817.2012.01224.x.

Hawkins, S. J., Sugden, H. E., Mieszkowska, N., Moore, P. J., Poloczanska, E., Leaper, R., et al. (2009). Consequences of climate-driven biodiversity changes for ecosystem functioning of North European rocky shores. Mar. Ecol. Prog. Ser. 396, 245–259. Available at: https://www.int-res.com/abstracts/meps/v396/p245-259/.

Helmuth, B., Broitman, B. R., Blanchette, C. A., Gilman, S., Halpin, P., Harley, C. D. G., et al. (2006). Mosaic patterns of thermal stress in the rocky intertidal zone: implications for climate change. Ecol. Monogr. 76, 461–479. doi:https://doi.org/10.1890/0012-9615(2006)076[0461:MPOTSI]2.0.CO;2.

Hothorn, T., Bretz, F., Westfall, P., Heiberger, R. M., Schuetzenmeister, A., Scheibe, S., et al. (2016). Package ‘multcomp.’ Simultaneous inference Gen. Parametr. Model. Proj. Stat. Comput. Vienna, Austria.

James, K., and Shears, N. T. (2016). Proliferation of the invasive kelp Undaria pinnatifida at aquaculture sites promotes spread to coastal reefs. Mar. Biol. 163, 34.

Jurgens, L. J., Rogers-Bennett, L., Raimondi, P. T., Schiebelhut, L. M., Dawson, M. N., Grosberg, R. K., et al. (2015). Patterns of Mass Mortality among Rocky Shore Invertebrates across 100 km of Northeastern Pacific Coastline. PLoS One 10, e0126280. Available at: https://doi.org/10.1371/journal.pone.0126280.

Lagos, V., and Cremades, J. (2004). Contribución al conocimiento de la biología del alga parda alimentaria Himanthalia elongata (Fucales, Phaeophyta) en las costas de Galicia. An. Biol. 0.

Lima, F. P., Gomes, F., Seabra, R., Wethey, D. S., Seabra, M. I., Cruz, T., et al. (2016). Loss of thermal refugia near equatorial range limits. Glob. Chang. Biol. 22, 254– 263. doi:10.1111/gcb.13115.

Lima, F. P., Queiroz, N., Ribeiro, P. A., Hawkins, S. J., and Santos, A. M. (2006). Recent changes in the distribution of a marine gastropod, Patella rustica Linnaeus, 1758, and their relationship to unusual climatic events. J. Biogeogr. 33, 812–822.

Lima, F. P., Queiroz, N., Ribeiro, P. A., Xavier, R., Hawkins, S. J., and Santos, A. M. (2009). First record of </> Halidrys siliquosa</i> on the Portuguese coast: counter-intuitive range expansion? Mar. Biodivers. Rec. 2, e1. doi:10.1017/S1755267208000018.

Lima, F. P., Ribeiro, P. A., Queiroz, N., Hawkins, S. J., and Santos, A. M. (2007). Do distributional shifts of northern and southern species of algae match the warming pattern? Glob. Chang. Biol. 13, 2592–2604.

Lima, F. P., and Wethey, D. S. (2012). Three decades of high-resolution coastal sea surface temperatures reveal more than warming. Nat. Commun. 3, 1–13.

Lowe, S., Browne, M., Boudjelas, S., and De Poorter, M. (2000). 100 of the world’s worst invasive alien species: a selection from the global invasive species database. Invasive Species Specialist Group Auckland.

Martínez, B., Arenas, F., Trilla, A., Viejo, R. M., and Carreño, F. (2015). Combining physiological threshold knowledge to species distribution models is key to improving forecasts of the future niche for macroalgae. Glob. Chang. Biol. 21, 1422–1433. doi:10.1111/gcb.12655.

McKnight, E., Spake, R., Bates, A., Smale, D. A., and Rius, M. (2021). Non-native species outperform natives in coastal marine ecosystems subjected to warming and freshening events. Glob. Ecol. Biogeogr. n/a. doi:https://doi.org/10.1111/geb.13318.

Mclean, M., Mouillot, D., Maureaud, A. A., Engelhard, G., Pinsky, M., and Correspondence, A. A. (2021). Disentangling tropicalization and deborealization in marine ecosystems under climate change. Curr. Biol. 31. doi:10.1016/j.cub.2021.08.034.

Meneghesso, C., Seabra, R., Broitman, B. R., Wethey, D. S., Burrows, M. T., Chan, B. K. K., et al. (2020). Remotely-sensed L4 SST underestimates the thermal fingerprint of coastal upwelling. Remote Sens. Environ. 237, 111588.

Mieszkowska, N., Kendall, M. A., Hawkins, S. J., Leaper, R., Williamson, P., Hardman-Mountford, N. J., et al. (2006). “Changes in the range of some common rocky shore species in Britain—a response to climate change?,” in Marine Biodiversity (Springer), 241–251.

Minchin, D., and Nunn, J. (2014). The invasive brown alga Undaria pinnatifida (Harvey) Suringar, 1873 (Laminariales: Alariaceae), spreads northwards in Europe. BioInvasions Rec. 3, 57–63. doi:10.3391/bir.2014.3.2.01.

Mineur, F., Arenas, F., Assis, J., Davies, A. J., Engelen, A. H., Fernandes, F., et al. (2015). European seaweeds under pressure: Consequences for communities and ecosystem functioning. J. Sea Res. 98, 91–108. doi:10.1016/j.seares.2014.11.004.

Miranda, P., Coelho, F. E. S., Tomé, A. R., Valente, M. A., Carvalho, A., Pires, C., et al. (2002). 20th century Portuguese climate and climate scenarios. Clim. Chang. Port. Scenar. Impacts Adapt. Meas. Proj. (Santos FD, Forbes K, Moita R, eds). Lisbon Gradiva Publ., 23–83.

Molenaar, F. J., Venekamp, L. A. H., and Breeman, A. M. (1996). Life-history regulation in the subtidal red algae Calliblepharis ciliata. Eur. J. Phycol. 31, 241– 247. doi:10.1080/09670269600651441.

Morelli, T. L., Barrows, C. W., Ramirez, A. R., Cartwright, J. M., Ackerly, D. D., Eaves, T. D., et al. (2020). Climate-change refugia: biodiversity in the slow lane. Front. Ecol. Environ. 18, 228–234. doi:https://doi.org/10.1002/fee.2189.

Mulas, M., and Bertocci, I. (2016). Devil’s tongue weed (Grateloupia turuturu Yamada) in northern Portugal: Passenger or driver of change in native biodiversity? Mar. Environ. Res. 118, 1–9. doi:https://doi.org/10.1016/j.marenvres.2016.04.007.

Norderhaug, K. M., Nedreaas, K., Huserbråten, M., and Moland, E. (2021). Depletion of coastal predatory fish sub-stocks coincided with the largest sea urchin grazing event observed in the NE Atlantic. Ambio 50, 163–173. doi:10.1007/s13280-020-01362-4.

Oksanen, J., Blanchet, F. G., Kindt, R., Legendre, P., Minchin, P. R., O’hara, R. B., et al. (2013). Package ‘vegan.’ Community Ecol. Packag. version 2, 1–295.

Pearson, G. A., Lago-Leston, A., and Mota, C. (2009). Frayed at the edges: selective pressure and adaptive response to abiotic stressors are mismatched in low diversity edge populations. J. Ecol. 97, 450–462. doi:10.1111/j.1365-2745.2009.01481.x.

Pereira, J., Monteiro, C., Santos, A. M., Seabra, R., Ribeiro, P., and Lima, F. P. (2021a). Intertidal Biodiversity along the Portuguese coast (2001-2002). GBIF.org. doi:10.15468/mbg5p3.

Pereira, J., Monteiro, C., Seabra, R., Lima, F., and Lima, F. P. (2022). Fine-scale abundance of rocky shore macroalgae species with distribution limits in NW Iberia in 2020/2021. ARPHA Prepr. 2, e80942.. doi:10.3897/ARPHAPREPRINTS.E80942.

Pereira, J., Monteiro, C., Seabra, R., Santos Múrias, A., and Lima, F. P. (2021b). Intertidal macroalgae species distribution along the North-western Iberian coast in 2020/2021. GBIF.org. doi:10.15468/247z4g.

Pereira, J., Ribeiro, P. A., Santos, A. M., Monteiro, C., Seabra, R., and Lima, F. P. (2021c). A comprehensive assessment of the intertidal biodiversity along the Portuguese coast in the early 2000s. Biodivers. Data J. 9. doi:10.3897/BDJ.9.e72961.

Petrov, S. (2021). Permutation hypothesis test in R. Exploring a powerful simulation… | by Serafim Petrov | Towards Data Science.

Piñeiro-Corbeira, C., Barreiro, R., and Cremades, J. (2016). Decadal changes in the distribution of common intertidal seaweeds in Galicia (NW Iberia). Mar. Environ. Res. 113, 106–115. doi:https://doi.org/10.1016/j.marenvres.2015.11.012.

Plaza, M., Cifuentes, A., and Ibáñez, E. (2008). In the search of new functional food ingredients from algae. Trends Food Sci. Technol. 19, 31–39.

Poloczanska, E. S., Burrows, M. T., Brown, C. J., García Molinos, J., Halpern, B. S., Hoegh-Guldberg, O., et al. (2016). Responses of Marine Organisms to Climate Change across Oceans. Front. Mar. Sci. 3, 62. Available at: https://www.frontiersin.org/article/10.3389/fmars.2016.00062.

Potter, K. A., Arthur Woods, H., and Pincebourde, S. (2013). Microclimatic challenges in global change biology. Glob. Chang. Biol. 19, 2932–2939. doi:10.1111/gcb.12257.

Poza, A. M., Gauna, M. C., Escobar, J. F., and Parodi, E. R. (2017). Heteromorphic phases of Leathesia marina (Ectocarpales, Ochrophyta) over time from northern Patagonia, Argentina. Phycologia 56, 579–589. doi:10.2216/16-117.1.

Provan, J., and Maggs, C. A. (2012). Unique genetic variation at a species’ rear edge is under threat from global climate change. Proc. R. Soc. B Biol. Sci. 279, 39–47. doi:10.1098/rspb.2011.0536.

R Core Team (2020). R: A Language and Environment for Statistical Computing.

Ramos, E., Guinda, X., Puente, A., de la Hoz, C.F., and Juanes, J. A. (2020). Changes in the distribution of intertidal macroalgae along a longitudinal gradient in the northern coast of Spain. Mar. Environ. Res. 157, 104930. doi:https://doi.org/10.1016/j.marenvres.2020.104930.

Rubal, M., Veiga, P., Cacabelos, E., Moreira, J., and Sousa-Pinto, I. (2013). Increasing sea surface temperature and range shifts of intertidal gastropods along the Iberian Peninsula. J. Sea Res. 77, 1–10. doi:https://doi.org/10.1016/j.seares.2012.12.003.

Rubal, M., Veiga, P., Maldonado, C., Torres, C., and Moreira, J. (2015). Population attributes and traits of Siphonaria pectinata (Mollusca: Siphonariidae) in range-edge and non range-edge populations at its Eastern Atlantic northern distribution boundary. J. Exp. Mar. Bio. Ecol. 471, 41–47. doi:10.1016/j.jembe.2015.05.015.

Rudis, B. (2020). Additional Themes, Theme Components and Utilities for “ggplot2” [R package hrbrthemes version 0.8.0].

Sanchez, I., and Fernandez, C. (2005). Impact of the invasive seaweed Sargassum muticum (Phaeophyta) on an intertidal macroalgal assemblage. J. Phycol. 41, 923– 930.

Schnute, J., Boers, N., and Haigh, R. (2004). PBS Mapping 2: User’s Guide. Can. Tech. Rep. Fish. Aquat. Sci. 2549, viii + 126 p.

Seabra, R., Varela, R., Santos, A. M., Gómez-Gesteira, M., Meneghesso, C., Wethey, D. S., et al. (2019). Reduced Nearshore Warming Associated With Eastern Boundary Upwelling Systems. Front. Mar. Sci. 6, 104. Available at: https://www.frontiersin.org/article/10.3389/fmars.2019.00104.

Seabra, R., Wethey, D. S., Santos, A. M., and Lima, F. P. (2011). Side matters: microhabitat influence on intertidal heat stress over a large geographical scale. J. Exp. Mar. Bio. Ecol. 400, 200–208.

Seabra, R., Wethey, D. S., Santos, A. M., and Lima, F. P. (2015). Understanding complex biogeographic responses to climate change. Sci. Rep. 5, 1–6.

Seitz, R. D., Wennhage, H., Bergström, U., Lipcius, R. N., and Ysebaert, T. (2014). Ecological value of coastal habitats for commercially and ecologically important species. ICES J. Mar. Sci. 71, 648–665. doi:10.1093/icesjms/fst152.

Silva, C. O., Lemos, M. F. L., Gaspar, R., Gonçalves, C., and Neto, J. M. (2021). The effects of the invasive seaweed Asparagopsis armata on native rock pool communities: Evidences from experimental exclusion. Ecol. Indic. 125, 107463. doi:https://doi.org/10.1016/j.ecolind.2021.107463.

Slowikowski, K., Schep, A., Hughes, S., Lukauskas, S., Irisson, J.-O., Kamvar, Z. N., et al. (2018). Package ggrepel. Autom. position non-overlapping text labels with ‘ggplot2.

Smale, D. A., Burrows, M. T., Moore, P., O’Connor, N., and Hawkins, S. J. (2013). Threats and knowledge gaps for ecosystem services provided by kelp forests: a northeast Atlantic perspective. Ecol. Evol. 3, 4016–4038. doi:https://doi.org/10.1002/ece3.774.

Sousa, M. C., Ribeiro, A., Des, M., Gomez-Gesteira, M., deCastro, M., and Dias, J. M. (2020). NW Iberian Peninsula coastal upwelling future weakening: Competition between wind intensification and surface heating. Sci. Total Environ. 703, 134808. doi:https://doi.org/10.1016/j.scitotenv.2019.134808.

South, P. M., Floerl, O., Forrest, B. M., and Thomsen, M. S. (2017). A review of three decades of research on the invasive kelp Undaria pinnatifida in Australasia: An assessment of its success, impacts and status as one of the world’s worst invaders. Mar. Environ. Res. 131, 243–257. doi:https://doi.org/10.1016/j.marenvres.2017.09.015.

Southward, A. J., Hawkins, S. J., and Burrows, M. T. (1995). Seventy years’ observations of changes in distribution and abundance of zooplankton and intertidal organisms in the western English Channel in relation to rising sea temperature. J. Therm. Biol. 20, 127–155. doi:10.1016/0306-4565(94)00043-I.

Stachowicz, J. J., Terwin, J. R., Whitlatch, R. B., and Osman, R. W. (2002). Linking climate change and biological invasions: Ocean warming facilitates nonindigenous species invasions. Proc. Natl. Acad. Sci. 99, 15497 LP – 15500. doi:10.1073/pnas.242437499.

Stæhr, P. A., Pedersen, M. F., Thomsen, M. S., Wernberg, T., and Krause-Jensen, D. (2000). Invasion of Sargassum muticum in Limfjorden (Denmark) and its possible impact on the indigenous macroalgal community. Mar. Ecol. Prog. Ser. 207, 79– 88.

Stuart, M. D., Hurd, C. L., and Brown, M. T. (1999). Effects of seasonal growth rate on morphological variation of Undaria pinnatifida (Alariaceae, Phaeophyceae). in Sixteenth International Seaweed Symposium, eds. J.M. Kain, M.T. Brown, and M. Lahaye (Dordrecht: Springer Netherlands), 191–199.

Teagle, H., and Smale, D. A. (2018). Climate-driven substitution of habitat-forming species leads to reduced biodiversity within a temperate marine community. Divers. Distrib. 24, 1367–1380. doi:https://doi.org/10.1111/ddi.12775.

Trivedi, M., Berry, P., Morecroft, M., and Dawson, T. (2008). Spatial Scale Affects Bioclimate Model Projections of Climate Change Impacts on Mountain Plants. Glob. Chang. Biol. 14, 1089–1103. doi:10.1111/j.1365-2486.2008.01553.x.

Ullah, H., Nagelkerken, I., Goldenberg, S. U., and Fordham, D. A. (2018). Climate change could drive marine food web collapse through altered trophic flows and cyanobacterial proliferation. PLOS Biol. 16, e2003446. Available at: https://doi.org/10.1371/journal.pbio.2003446.

Van den Hoek, C., and Donze, M. (1967). Algal phytogeography of the European Atlantic coasts. Blumea Biodiversity, Evol. Biogeogr. Plants 15, 63–89.

Veiga, P., Torres, A. C., Rubal, M., Troncoso, J., and Sousa-Pinto, I. (2014). The invasive kelp </> Undaria pinnatifida</i> (Laminariales, Ochrophyta) along the north coast of Portugal: Distribution model versus field observations. Mar. Pollut. Bull. 84, 363–365.

Viana, I. G., Bode, A., and Fernández, C. (2014). Growth and production of new recruits and adult individuals of Ascophyllum nodosum in a non-harvested population at its southern limit (Galicia, NW Spain). Mar. Biol. 161, 2885–2895. doi:10.1007/s00227-014-2553-0.

Vieira, R., Pereira, R., Arenas, F., Araujo, R., and Pinto, I. S. (2010). Espécies intertidais características da costa norte de Portugal., ed. SerSilito.

Wethey, D. S., Woodin, S. A., Hilbish, T. J., Jones, S. J., Lima, F. P., and Brannock, P. M. (2011). Response of intertidal populations to climate: effects of extreme events versus long term change. J. Exp. Mar. Bio. Ecol. 400, 132–144.

Whittick, A. (1983). Spatial and temporal distributions of dominant epiphytes on the stipes of Laminaria hyperborea (Gunn.) Fosl. (Phaeophyta:Laminariales) in S.E. Scotland. J. Exp. Mar. Bio. Ecol. 73, 1–10. doi:https://doi.org/10.1016/0022-0981(83)90002-3.

Wickham, H. (2012). reshape2: Flexibly reshape data: a reboot of the reshape package. R Packag. version 1.

Wickham, H., Chang, W., Henry, L., Pedersen, T. L., Takahashi, K., Wilke, C., et al. (2016). ggplot2: Create elegant data visualisations using the grammar of graphics. R Packag. version 2.

Williams, S. L., and Smith, J. E. (2007). A global review of the distribution, taxonomy, and impacts of introduced seaweeds. Annu. Rev. Ecol. Evol. Syst. 38, 327–359.

Yesson, C., Bush, L. E., Davies, A. J., Maggs, C. A., and Brodie, J. (2015). Large brown seaweeds of the British Isles: Evidence of changes in abundance over four decades. Estuar. Coast. Shelf Sci. 155, 167–175. doi:https://doi.org/10.1016/j.ecss.2015.01.008.

