## Supplementary material for "Fine-scale survey of intertidal macroalgae reveals recent changes in a cold-water biogeographic stronghold"

### *Suplementary material:*

**Supplementary Table 1-** List of species surveyed, scientific name from the World Register of Marine Species (WoRMS), temperature affinity, and geographical origin.

| Scientific name | Temperature affinity | Geographical origin |
|---|---|---|
| *Ascophyllum nodosum* (Linnaeus) Le Jolis, 1863 | Cold-water species | Native species |
| *Asparagopsis armata* Harvey, 1855 | Warm-water species | Invasive species |
| *Calliblepharis ciliata* (Hudson) Kützing, 1843 | Cold-water species | Native species |
| *Chondrus crispus* Stackhouse, 1797 | Cold-water species | Native species |
| *Codium adhaerens* C.Agardh, 1822 | Warm-water species | Native species |
| *Delesseria sanguinea* (Hudson) J.V.Lamouroux, 1813 | Cold-water species | Native species |
| *Desmarestia ligulata* (Stackhouse) J.V.Lamouroux, 1813 | Cold-water species | Native species |
| *Dictyopteris polypodioides* (A.P.De Candolle) J.V.Lamouroux, 1809 | Warm-water species | Native species |
| *Dilsea carnosa* (Schmidel) Kuntze, 1898 | Cold-water species | Native species |
| *Dumontia contorta* (S.G.Gmelin) Ruprecht, 1850 | Cold-water species | Native species |
| *Fucus serratus* Linnaeus, 1753 | Cold-water species | Native species |
| *Fucus spiralis* Linnaeus, 1753 | Cold-water species | Native species |
| *Gelidium corneum* (Hudson) J.V.Lamouroux, 1813 | Warm-water species | Native species |
| *Grateloupia turuturu* Yamada, 1941 | Neutral species | Invasive species |
| *Halidrys siliquosa* (Linnaeus) Lyngbye, 1819 | Cold-water species | Native species |
| *Halopithys incurva* (Hudson) Batters, 1902 | Warm-water species | Native species |
| *Himanthalia elongata* (Linnaeus) S.F.Gray, 1821 | Cold-water species | Native species |
| *Hypnea musciformis* (Wulfen) J.V.Lamouroux, 1813 | Warm-water species | Native species |
| *Laminaria hyperborea* (Gunnerus) Foslie, 1884 | Cold-water species | Native species |
| *Laminaria ochroleuca* Bachelot de la Pylaie, 1824 | Warm-water species | Native species |
| *Leathesia marina* (Lyngbye) Decaisne, 1842 | Cold-water species | Native species |
| *Padina pavonica* (Linnaeus) Thivy, 1960 | Warm-water species | Native species |
| *Palmaria palmata* (Linnaeus) F.Weber & D.Mohr, 1805 | Cold-water species | Native species |
| *Pelvetia canaliculata* (Linnaeus) Decaisne & Thuret, 1845 | Cold-water species | Native species |

| | | |
|---|---|---|
| *Petalonia fascia* (O.F.Müller) Kuntze, 1898 | Neutral species | Native species |
| *Phycodrys rubens* (Linnaeus) Batters, 1902 | Cold-water species | Native species |
| *Phyllophora crispa* (Hudson) P.S.Dixon, 1964 | Neutral species | Native species |
| *Treptacantha baccata* (S.G.Gmelin) Orellana & Sansón, 2019 | Neutral species | Native species |
| *Saccharina latissima* (Linnaeus) C. E. Lane, C. Mayes, Druehl & G. W. Saunders, 2006 | Cold-water species | Native species |
| *Saccorhiza polyschides* (Lightfoot) Batters, 1902 | Cold-water species | Native species |
| *Sargassum flavifolium* Kützing, 1849 | Warm-water species | Native species |
| *Sargassum muticum* (Yendo) Fensholt, 1955 | Neutral species | Invasive species |
| *Undaria pinnatifida* (Harvey) Suringar, 1873 | Cold-water species | Invasive species |
| *Valonia utricularis* (Roth) C.Agardh, 1823 | Warm-water species | Native species |

**Supplementary Table 1** – Rocky shore sites surveyed, location ID, Location name, coordinates, date and astronomical low tide of the day of the survey. The coordinates were obtained from GoogleMaps imagery. Locations are listed from North to South. Sites with an * were also surveyed in 2001-2005 (Pereira et al., 2021a).

| ID | Location | Latitude | Longitude | Date | Astronomical low tide height (m below mean sea water level) |
|---|---|---|---|---|---|
| 1 | Cabo Touriñán* | 43.04423 | -9.2881 | 24/05/2021 | -1.52171 |
| 2 | Corveiro | 42.90442 | -9.26077 | 26/05/2021 | -1.68245 |
| 3 | Quenxe | 42.9365 | -9.18958 | 26/05/2021 | -1.67836 |
| 4 | Ximprón | 42.79679 | -9.14016 | 25/05/2021 | -1.63124 |
| 5 | Punta Outeiriño | 42.74564 | -9.07681 | 25/05/2021 | -1.62846 |
| 6 | Corrubedo* | 42.57665 | -9.08985 | 26/06/2021 | -1.44851 |
| 7 | O Touro | 42.54606 | -8.98397 | 26/06/2021 | -1.45043 |
| 8 | Prado | 42.15921 | -8.8194 | 27/05/2021 | -1.6242 |
| 9 | Faro Vello de Silleiro | 42.11185 | -8.89945 | 27/05/2021 | -1.61985 |
| 10 | Oia | 42.00199 | -8.8777 | 28/05/2021 | -1.49061 |
| 11 | Fedorento | 41.91017 | -8.87801 | 28/05/2021 | -1.48237 |
| 12 | Moledo* | 41.83815 | -8.87491 | 19/10/2020 | -1.69232 |
| 12 | Moledo* | 41.83908 | -8.87529 | 25/06/2021 | -1.42987 |
| 13 | Vila Praia de Âncora* | 41.8194 | -8.87205 | 17/12/2020 | -1.45051 |

| | | | | | |
|---|---|---|---|---|---|
| 14 | Afife* | 41.78439 | -8.87168 | 17/12/2020 | -1.68671 |
| 14 | Afife* | 41.78072 | -8.87014 | 19/10/2020 | -1.44752 |
| 15 | Montedor* | 41.74292 | -8.87591 | 29/01/2021 | -1.36943 |
| 16 | Forte da Vigia* | 41.69959 | -8.85507 | 16/11/2020 | -1.71516 |
| 17 | Praia Norte* | 41.69983 | -8.85472 | 16/11/2020 | -1.71516 |
| 18 | Amorosa* | 41.6429 | -8.82338 | 12/01/2021 | -1.38846 |
| 19 | Foz do Neiva | 41.61095 | -8.80893 | 16/12/2020 | -1.54521 |
| 20 | Rio de Moinhos | 41.57362 | -8.79846 | 16/12/2020 | -1.54521 |
| 21 | Apúlia | 41.48267 | -8.77886 | 17/11/2020 | -1.62315 |
| 22 | Santo André | 41.41663 | -8.78827 | 15/01/2021 | -1.42303 |
| 23 | Verde | 41.38542 | -8.77433 | 15/01/2021 | -1.42238 |
| 24 | Carvalhido | 41.38149 | -8.7715 | 30/03/2021 | -1.79394 |
| 25 | Caxinas | 41.3622 | -8.76045 | 13/01/2021 | -1.46149 |
| 26 | Forte de São João | 41.34108 | -8.75073 | 13/01/2021 | -1.46137 |
| 27 | Areia | 41.33355 | -8.73993 | 14/01/2021 | -1.48803 |
| 28 | Mindelo* | 41.31052 | -8.74136 | 14/01/2021 | -1.48853 |
| 29 | Facho* | 41.29241 | -8.73419 | 15/12/2020 | -1.57976 |
| 30 | Sampaio | 41.27956 | -8.72914 | 15/12/2020 | -1.57976 |
| 31 | Labruge* | 41.27309 | -8.729 | 16/01/2021 | -1.3163 |
| 32 | Angeiras (Maelas) | 41.26615 | -8.72829 | 31/03/2021 | -1.68339 |
| 33 | Angeiras (Praia dos Barcos)* | 41.2651 | -8.72818 | 16/01/2021 | -1.3163 |
| 34 | Praia Central | 41.26187 | -8.72686 | 31/01/2021 | -1.4829 |
| 35 | Funtão | 41.26041 | -8.72494 | 15/11/2020 | -1.65946 |
| 36 | Pedras do Corgo | 41.24931 | -8.72591 | 15/11/2020 | -1.65947 |
| 37 | Pedras da Agudela | 41.24163 | -8.72795 | 14/11/2020 | -1.54667 |
| 38 | Memória | 41.23528 | -8.72433 | 17/10/2020 | -1.71118 |
| 39 | Cabo do Mundo* | 41.22115 | -8.71577 | 17/10/2020 | -1.71134 |
| 40 | Boa Nova | 41.20458 | -8.71553 | 16/10/2020 | -1.59479 |

| | | | | | |
|---|---|---|---|---|---|
| 41 | Leça (Piscina das Marés) | 41.19231 | -8.70742 | 16/10/2020 | -1.59497 |
| 42 | Castelo do Queijo | 41.16746 | -8.69016 | 15/10/2020 | -1.37579 |
| 42 | Castelo do Queijo | 41.16722 | -8.6902 | 23/06/2021 | -1.34778 |
| 43 | Homem do Leme* | 41.15903 | -8.68538 | 14/12/2020 | -1.54155 |
| 43 | Homem do Leme* | 41.15903 | -8.68538 | 14/02/2021 | -1.37303 |
| 44 | Salgueiros | 41.12148 | -8.66652 | 18/11/2020 | -1.45485 |
| 45 | Valadares* | 41.08964 | -8.657 | 18/11/2020 | -1.45516 |
| 46 | Senhor da Pedra* | 41.06894 | -8.65836 | 18/10/2020 | -1.75618 |
| 46 | Senhor da Pedra* | 41.06846 | -8.65848 | 24/06/2021 | -1.4027 |
| 47 | Aguda* | 41.04554 | -8.65282 | 18/10/2020 | -1.75629 |
| 47 | Aguda* | 41.04613 | -8.65325 | 24/06/2021 | -1.40298 |
| 48 | Buarcos* | 40.17751 | -8.90354 | 03/03/2021 | -1.46205 |
| 49 | Nazaré* | 39.60384 | -9.08041 | 01/03/2021 | -1.62904 |
| 50 | São Martinho do Porto* | 39.51151 | -9.14207 | 26/07/2021 | -1.27679 |
| 51 | Baleal* | 39.37586 | -9.33981 | 02/03/2021 | -1.57911 |
| 52 | Papôa* | 39.37344 | -9.37773 | 02/03/2021 | -1.57954 |

**Supplementary Table 3** - Permutational Multivariate Analysis of Variance Using Distance Matrices, based on Bray-Curtis dissimilarity, to detect differences between the regions Galicia, northern Portugal and central Portugal. The test used 999 random permutations.

| **Permutational Multivariate Analysis of Variance Using Distance Matrices** | | | | | | | |
|---|---|---|---|---|---|---|---|
| | Df | Sums of Squares | Mean Squares | F. Model | $R^2$ | Pr (>F) | sig |
| Regions | 2 | 1.773 | 0.8865 | 10.021 | 0.29455 | 0.001 | *** |
| Residuals | 48 | 4.2462 | 0.08846 | | 0.70545 | | |
| Total | 50 | 6.0192 | | | 1 | | |

**Supplementary Table 4** - Summary of Pairwise comparisons results, based on Bray-Curtis dissimilarity, between the geographical groups: Galicia (G), northern Portugal (NPT) and central Portugal (CPT). Test with 999 random permutations

| PARWISE PERMANOVA | | | | | | | |
|---|---|---|---|---|---|---|---|
| pairs | Df | Sums of Squares | F. Model | $R^2$ | p-value | p-adjusted | sig |
| G vs NPT | 1 | 1.0728786 | 12.590294 | 0.2186183 | 0.001 | 0.003 | * |
| G vs CPT | 1 | 0.3913105 | 4.693616 | 0.2652717 | 0.002 | 0.006 | * |
| NPT vs CPT | 1 | 0.8531891 | 9.07139 | 0.1927156 | 0.001 | 0.003 | * |

**Supplementary Table 5**- List of species that characterise regions and groups of regions - Galicia (G), northern Portugal (NPT) and central Portugal (CPT), based on Multilevel pattern analysis. Indicator values (IndVal) of species with highest proportion of specificity (A) and fidelity (B) are described by region and groups of regions. p-values are given by permutational analysis (n=9999).

| Group G #sps. 4 | | | | | |
|---|---|---|---|---|---|
| | A | B | stat | p-value | |
| *Undaria pinnatifida* | 0.9305 | 1 | 0.965 | 0.0001 | *** |
| *Himanthalia elongata* | 0.83721 | 0.27273 | 0.478 | 0.0864 | . |
| *Halidrys siliquosa* | 1 | 0.09091 | 0.302 | 0.2993 | |
| *Padina pavonica* | 1 | 0.09091 | 0.302 | 0.2993 | |
| Group NPT #sps. 5 | | | | | |
| | A | B | stat | p-value | |
| *Laminaria hyperborea* | 1 | 0.33333 | 0.577 | 0.102 | |
| *Palmaria palmata* | 1 | 0.05556 | 0.236 | 1 | |
| *Ascophyllum nodosum* | 1 | 0.02778 | 0.167 | 1 | |
| *Fucus serratus* | 1 | 0.02778 | 0.167 | 1 | |
| *Saccharina latissima* | 1 | 0.02778 | 0.167 | 1 | |
| Group CPT #sps. 3 | | | | | |
| | A | B | stat | p-value | |
| *Leathesia marina* | 1 | 0.5 | 0.707 | 0.0044 | ** |
| *Codium adhaerens* | 0.6735 | 0.5 | 0.58 | 0.0429 | * |
| *Sargassum flavifolium* | 1 | 0.25 | 0.5 | 0.0757 | . |
| Group G+NPT #sps. 6 | | | | | |
| | A | B | stat | p-value | |
| *Sargassum muticum* | 1 | 0.9149 | 0.957 | 0.0008 | *** |
| *Grateloupia turuturu* | 1 | 0.8085 | 0.899 | 0.0036 | ** |
| *Laminaria ochroleuca* | 1 | 0.5745 | 0.758 | 0.0764 | . |
| *Dilsea carnosa* | 1 | 0.1702 | 0.413 | 0.7301 | |
| *Pelvetia canaliculata* | 1 | 0.1702 | 0.413 | 0.737 | |
| *Desmarestia ligulata* | 1 | 0.1064 | 0.326 | 1 | |
| Group CPT+G #sps. 2 | | | | | |
| | A | B | stat | p-value | |
| *Asparagopsis armata* | 0.9235 | 0.8 | 0.86 | 0.0003 | *** |
| *Gelidium corneum* | 0.8251 | 0.6667 | 0.742 | 0.0479 | * |

| Group CPT+NPT #sps. 1 |  |  |  |  |
|---|---|---|---|---|
| | A | B | stat | p-value |
| *Calliblepharis ciliata* | 1 | 0.3 | 0.548 | 0.197 |

**Supplementary Table 6** - Krustal-Wallis sum test on the averaged Species Temperature Index (STI) by region on north-western Iberia.

| | **Kruskal-Wallis sum test** | | |
|---|---|---|---|
| | chi-squared | df | p-value |
| CTI | 8.8697 | 2 | 0.01186 |

**Supplementary Table 7** - Summary of Pairwise comparisons results, using Wilcoxon rank sum exact test, between the Community Temperature Index of the regions: Galicia (G), northern Portugal (NPT) and central Portugal (CPT).

| **Pairwise comparisons using Wilcoxon rank sum exact test** | | |
|---|---|---|
| | CPT | G |
| G | 1 | - |
| NPT | 0.5854 | 0.0089 |

**Supplementary Table 8** - Permutational Multivariate Analysis of Variance Using Distance Matrices, based on Bray-Curtis dissimilarity, to detect differences between the two time periods - historical (2001-2005) and current (2020-2021). The test used 999 random permutations.

| **Permutational Multivariate Analysis of Variance Using Distance Matrices** | | | | | | | |
|---|---|---|---|---|---|---|---|
| | Df | Sums of Squares | Mean Squares | F. Model | $R^2$ | Pr (>F) | sig |
| Time period | 1 | 0.753 | 0.75298 | 4.5558 | 0.09383 | 0.001 | *** |
| Residuals | 44 | 7.2723 | 0.16528 | | 0.90617 | | |
| Total | 45 | 8.0253 | | | 1 | | |

**Supplementary Table 8** - List of species that characterise historical (2001-2005) and current (2020-2021) sampling events, based on Multilevel pattern analysis. Indicator values (IndVal) of species with highest proportion of specificity (A) and fidelity (B) are described for each group. p-values are given by permutational analysis (n=9999).

| Historical group #sps. 8 |  |  |  |  |  |
|---|---|---|---|---|---|
| | A | B | stat | p-value | |
| *Himanthalia elongata* | 0.86111 | 0.3913 | 0.58 | 0.0123 | * |
| *Palmaria palmata* | 0.83333 | 0.30435 | 0.504 | 0.0399 | * |
| *Padina pavonica* | 1 | 0.13043 | 0.361 | 0.2324 | |
| *Delesseria sanguinea* | 1 | 0.08696 | 0.295 | 0.4873 | |
| *Dumontia contorta* | 1 | 0.04348 | 0.209 | 1 | |
| *Petalonia fascia* | 1 | 0.04348 | 0.209 | 1 | |

| | | | | | |
|---|---|---|---|---|---|
| *Phycodrys rubens* | 1 | 0.04348 | 0.209 | 1 | |
| *Phyllophora crispa* | 1 | 0.04348 | 0.209 | 1 | |
| Present-day group #sps. 5 | | | | | |
| | A | B | stat | p-value | |
| *Grateloupia turuturu* | 0.94118 | 0.73913 | 0.834 | 0.0001 | *** |
| *Sargassum muticum* | 0.75556 | 0.73913 | 0.747 | 0.002 | ** |
| *Calliblepharis ciliata* | 0.78947 | 0.26087 | 0.454 | 0.1643 | |
| *Undaria pinnatifida* | 1 | 0.13043 | 0.361 | 0.23 | |
| *Sargassum flavifolium* | 1 | 0.04348 | 0.209 | 1 | |

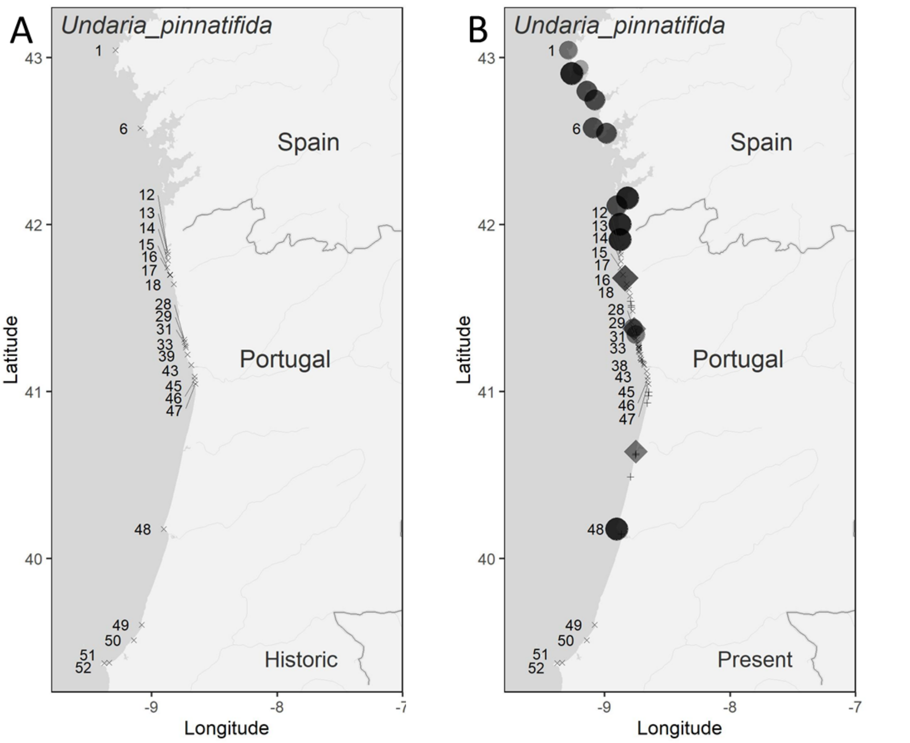

**Supplementary Figure 1** - Distribution maps of *Undaria pinnatifida* derived from historical (A) and present-day data (B). Rhombuses indicate the presence of the species in artificial sites and + indicate artificial sites in which the species was absent.

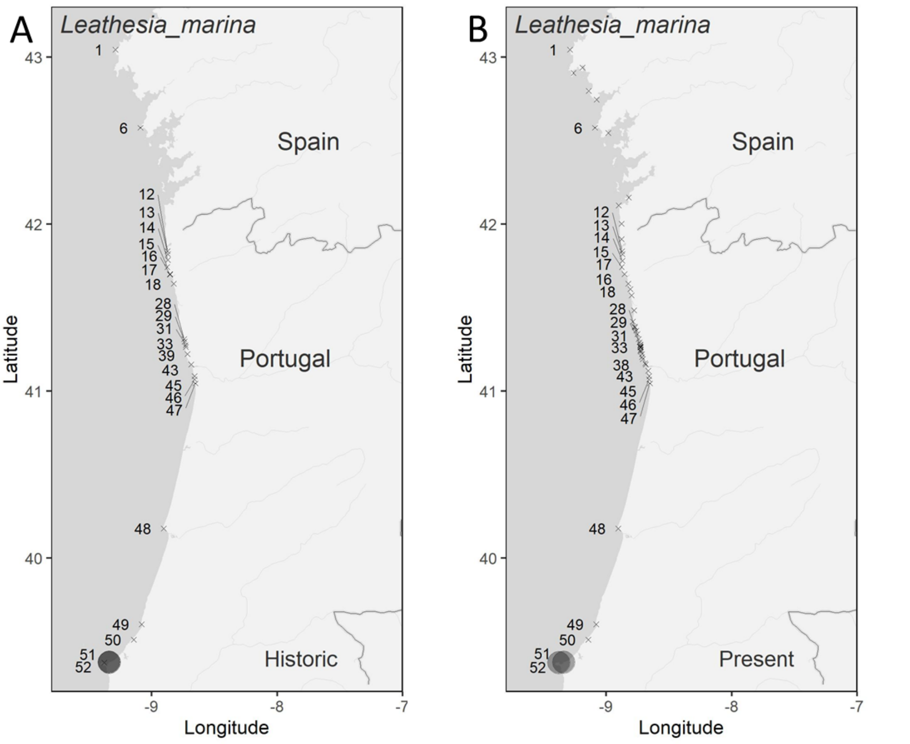

**Supplementary Figure 2** - Distribution maps of *Leathesia marina* derived from historical (A) and present-day data (B).

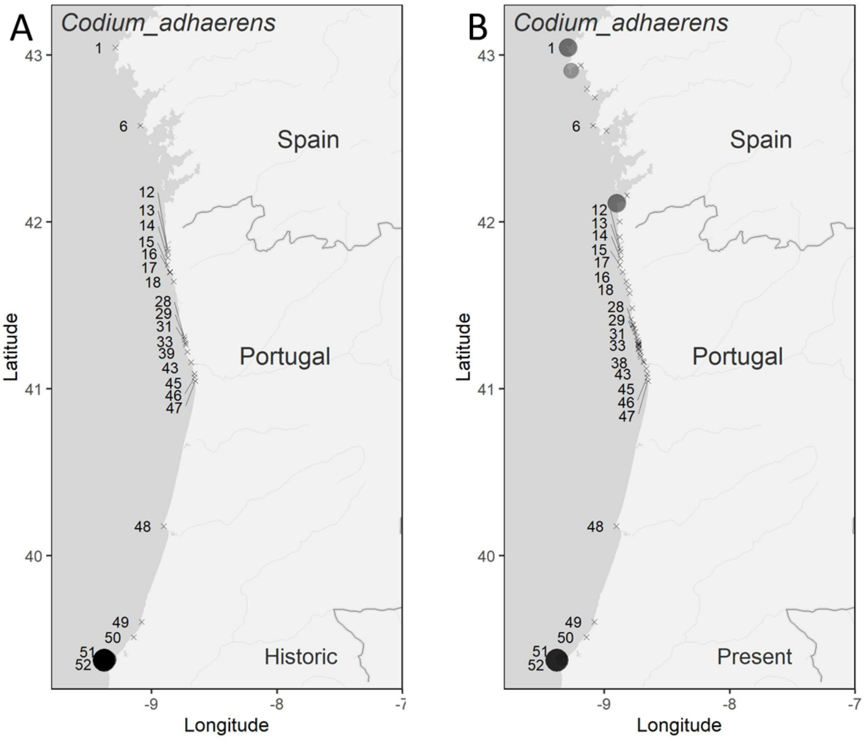

**Supplementary Figure 3** - Distribution maps of *Codium adhaerens* derived from historical (A) and present-day data (B).

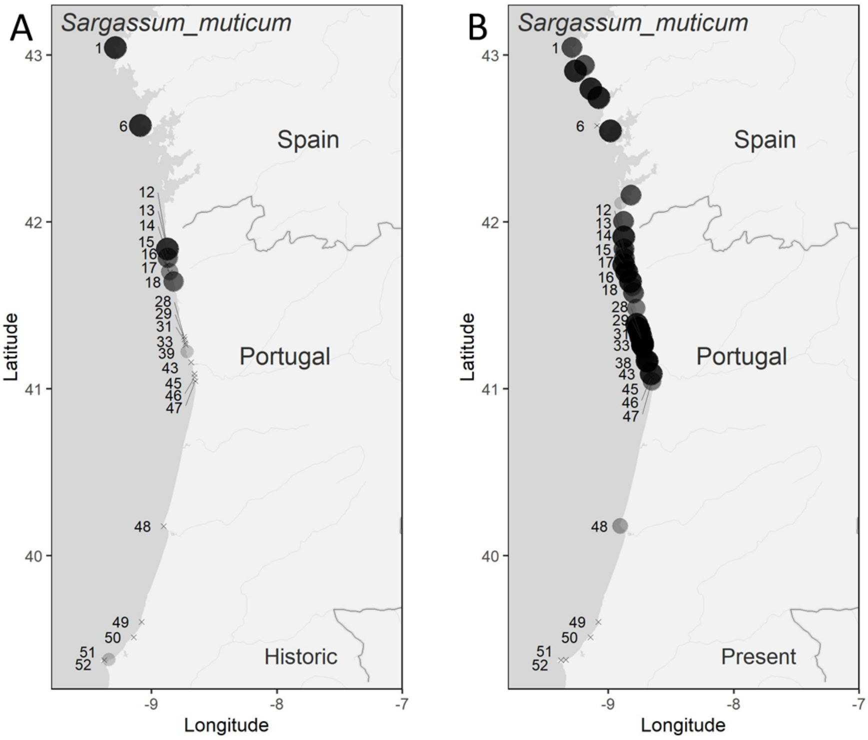

**Supplementary Figure 1** - Distribution maps of *Sargassum muticum* derived from historical (A) and present-day data (B).

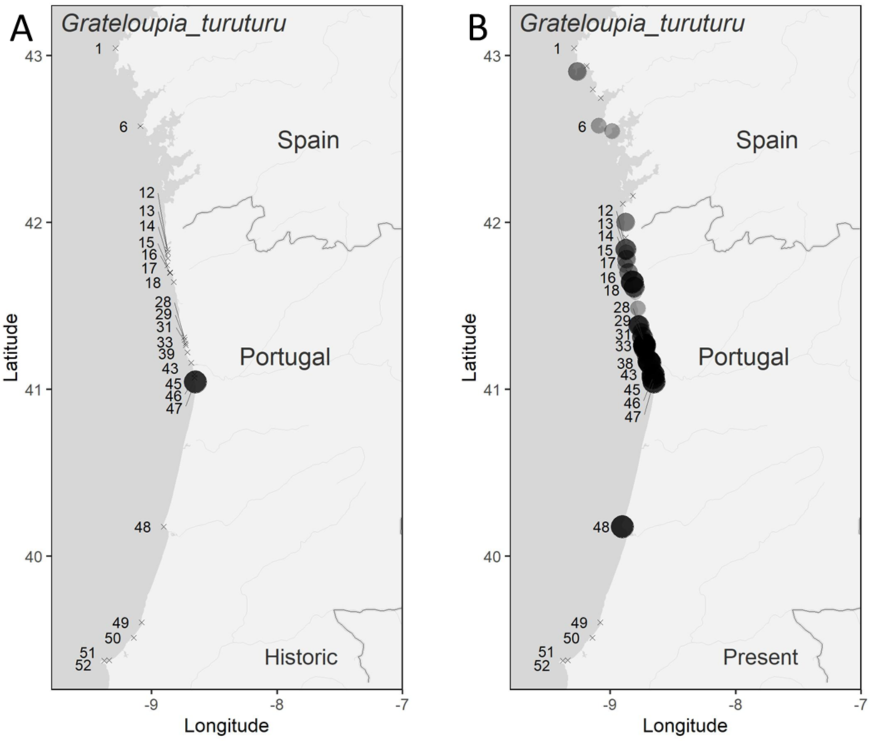

**Supplementary Figure 5** - Distribution maps of *Grateloupia turuturu* derived from historical (A) and present-day data (B).

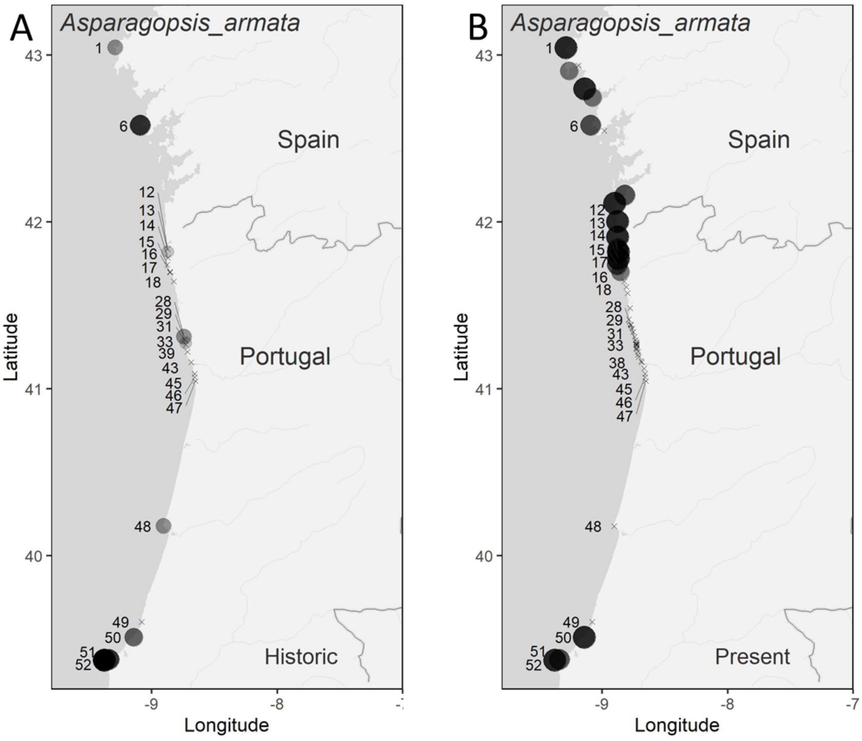

**Supplementary Figure 2** - Distribution maps of *Asparagopsis armata* derived from historical (A) and present-day data (B).

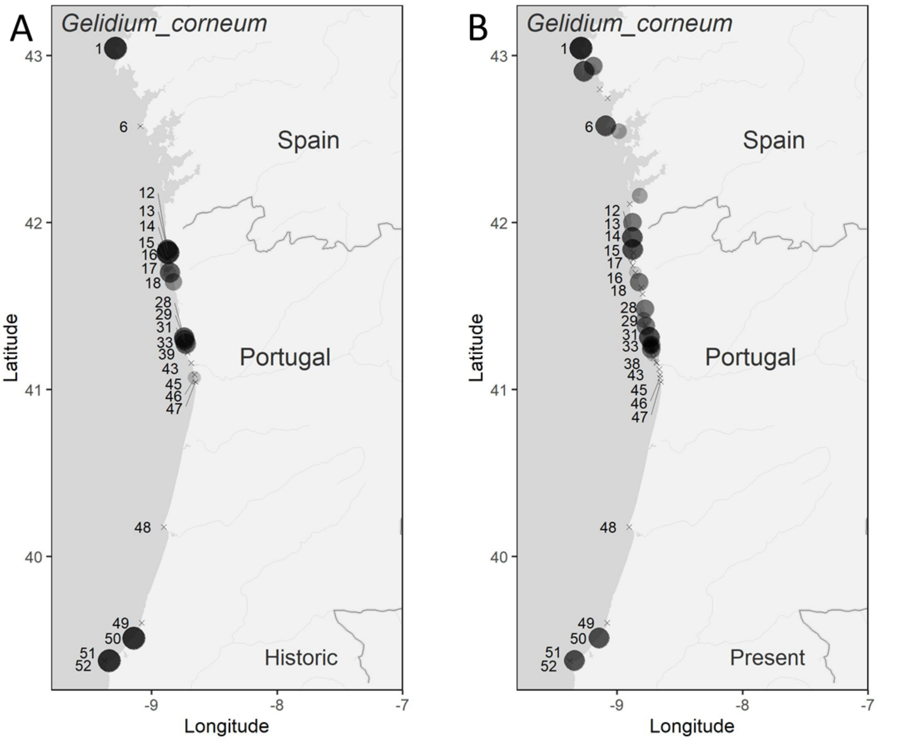

**Supplementary Figure 3** - Distribution maps of *Gelidium corneum* derived from historical (A) and present-day data (B).

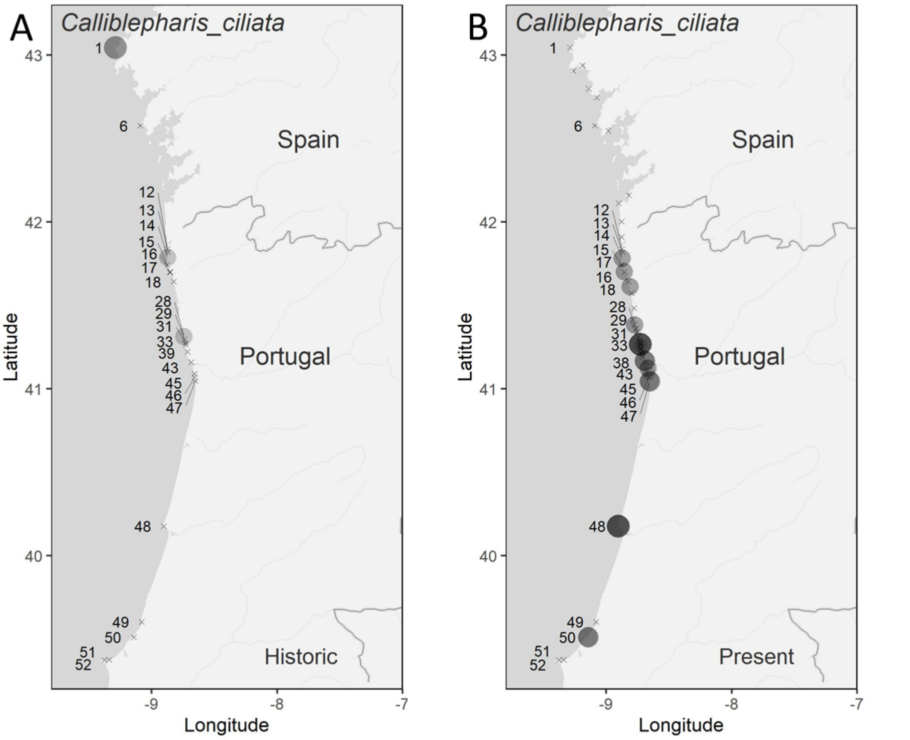

**Supplementary Figure 8** - Distribution maps of *Calliblepharis ciliata* derived from historical (A) and present-day data (B).

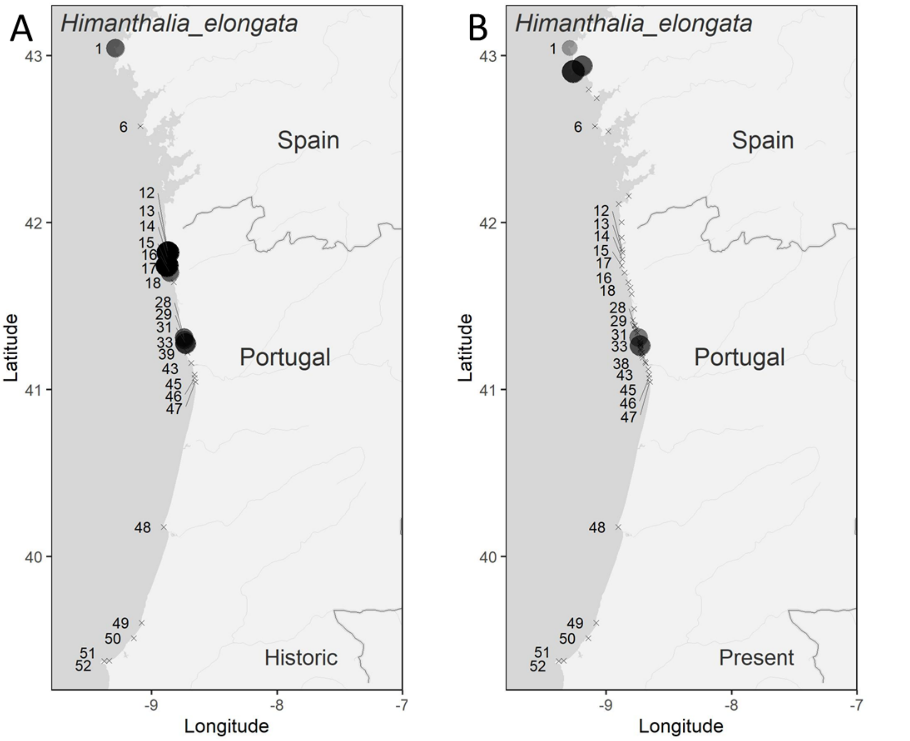

**Supplementary Figure 9** - Distribution maps of *Himanthalia elongata* derived from historical (A) and present-day data (B).

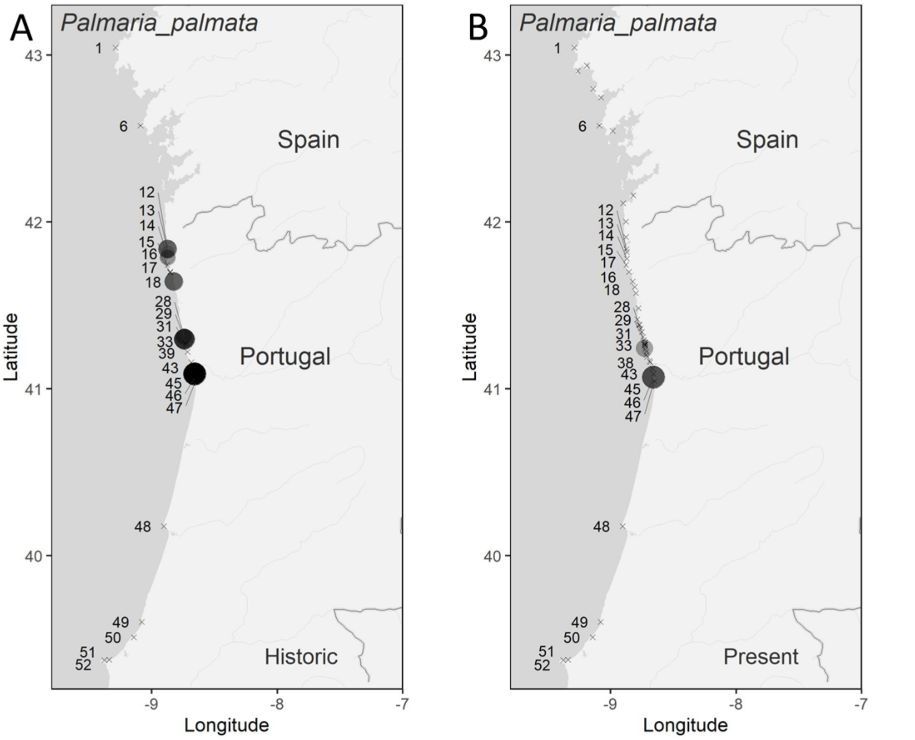

**Supplementary Figure 4** - Distribution maps of *Palmaria palmata* derived from historical (A) and present-day data (B).

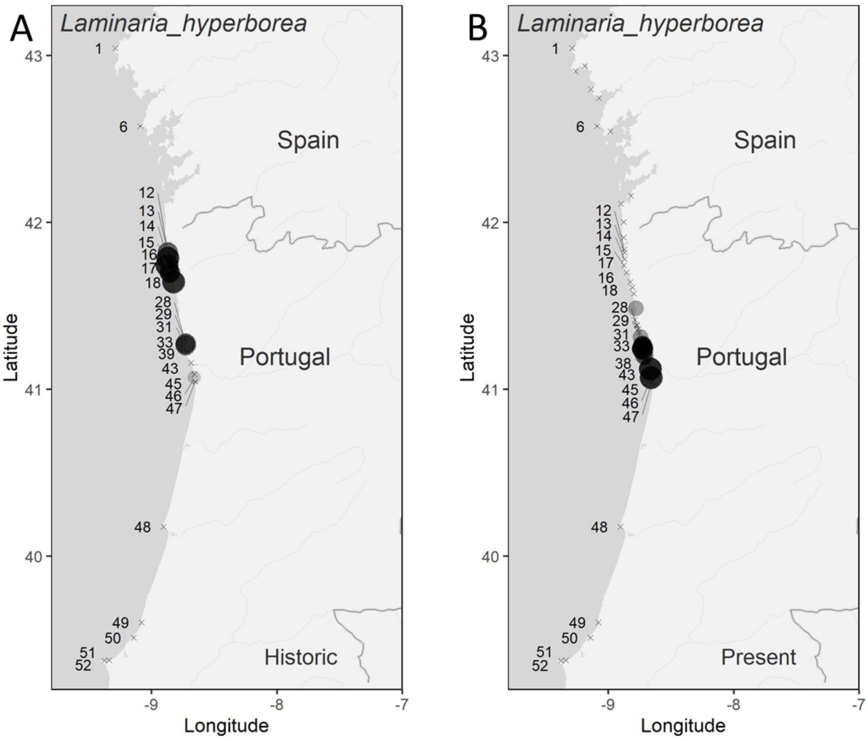

**Supplementary Figure 11** - Distribution maps of *Laminaria hyperborea* derived from historical (A) and present-day data (B).

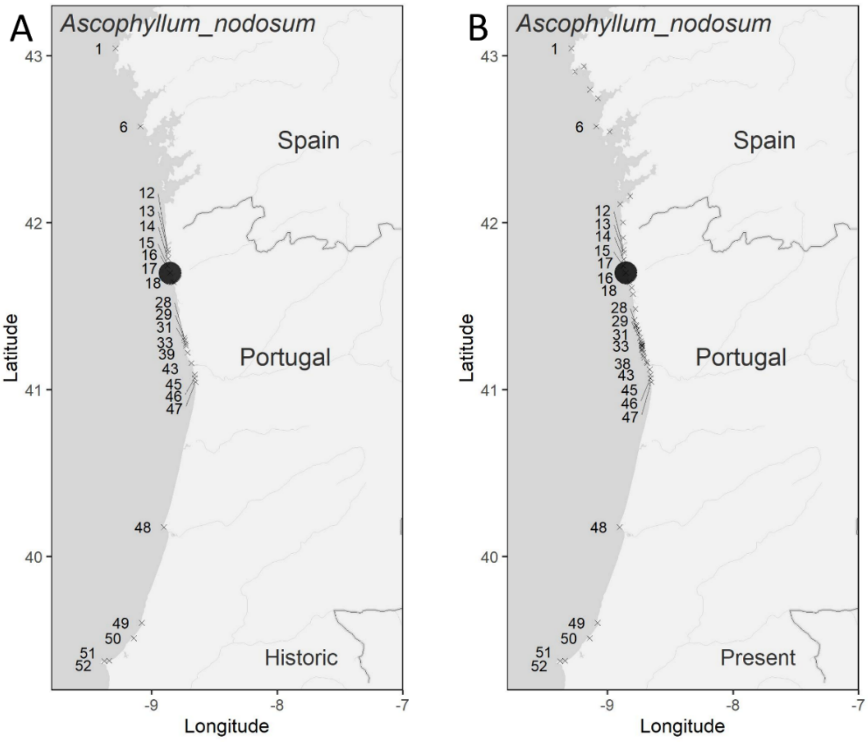

**Supplementary Figure 12**- Distribution maps of *Ascophyllum nodosum* derived from historical (A) and present-day data (B).

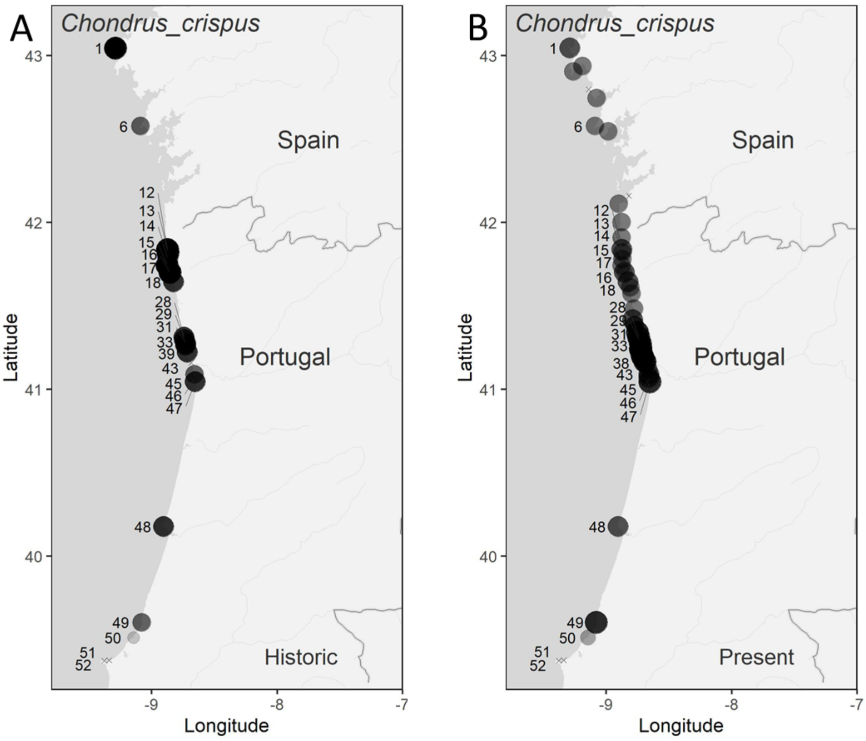

**Supplementary Figure 13**- Distribution maps of *Chondrus crispus* derived from historical (A) and present-day data (B).

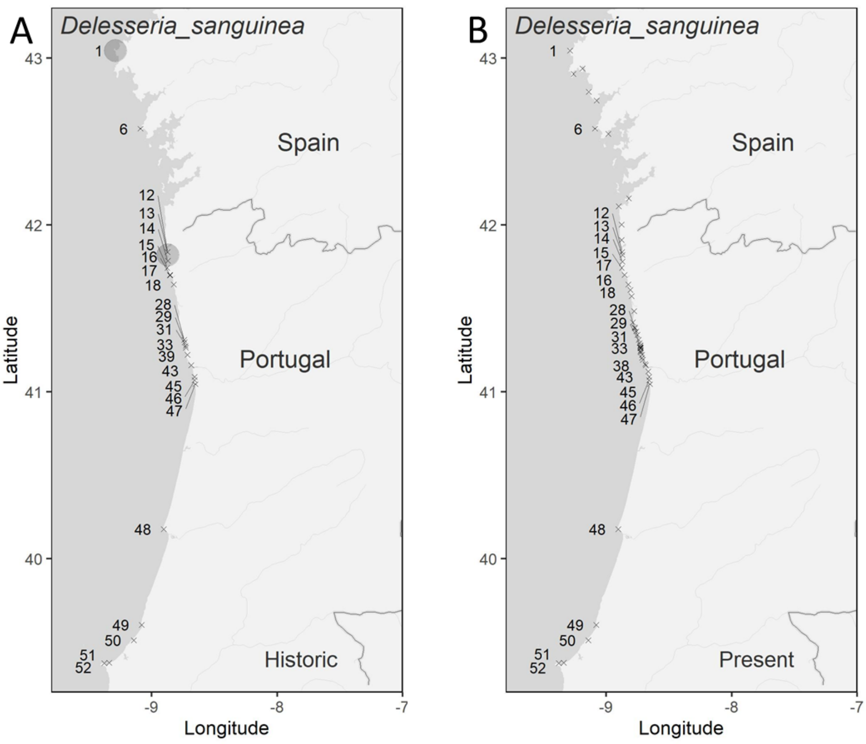

**Supplementary Figure 14** - Distribution maps of *Delesseria sanguinea* derived from historical (A) and present-day data (B).

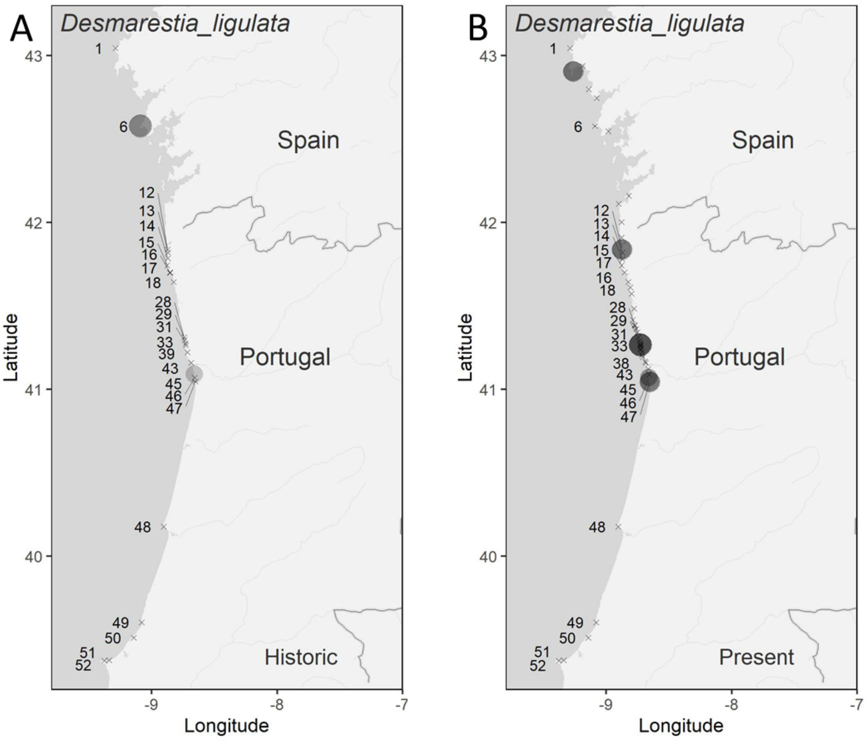

**Supplementary Figure 15** - Distribution maps of *Desmarestia ligulata* derived from historical (A) and present-day data (B).

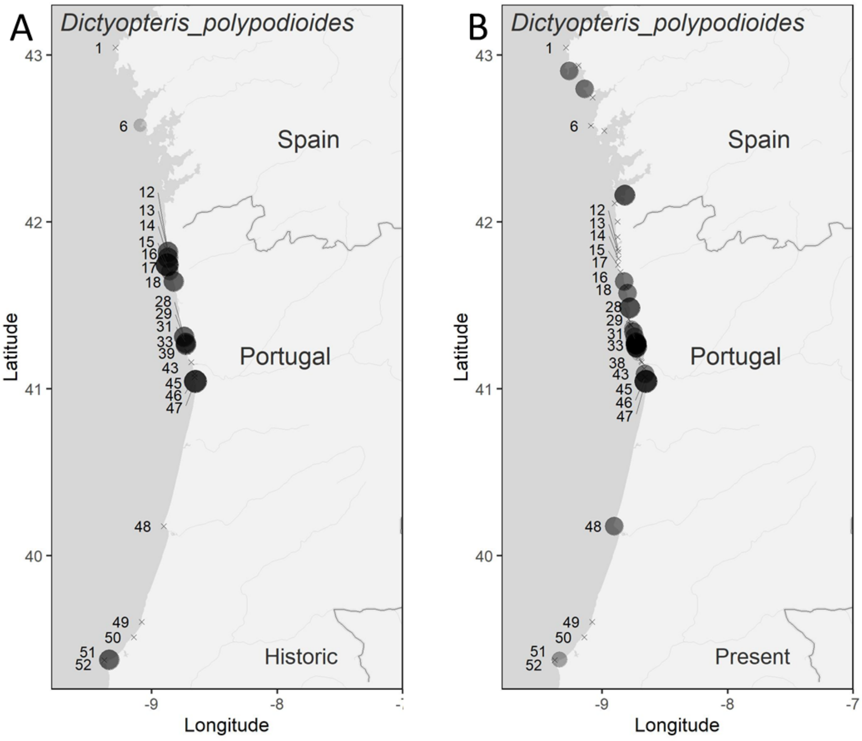

**Supplementary Figure 16** - Distribution maps of *Dictyopteris polypodioides* derived from historical (A) and present-day data (B).

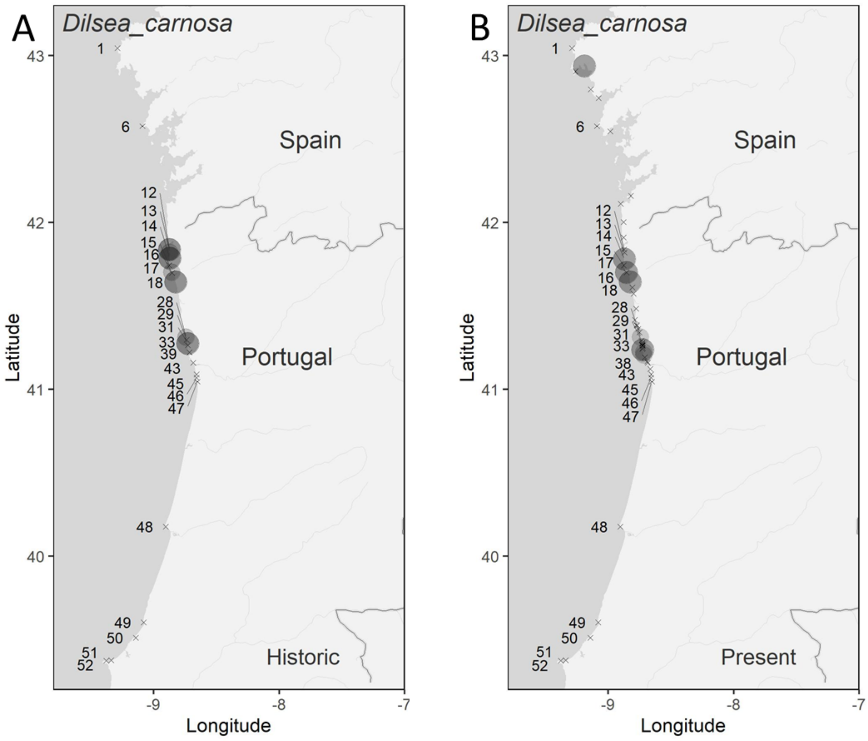

**Supplementary Figure 175** - Distribution maps of *Dilsea carnosa* derived from historical (A) and present-day data (B).

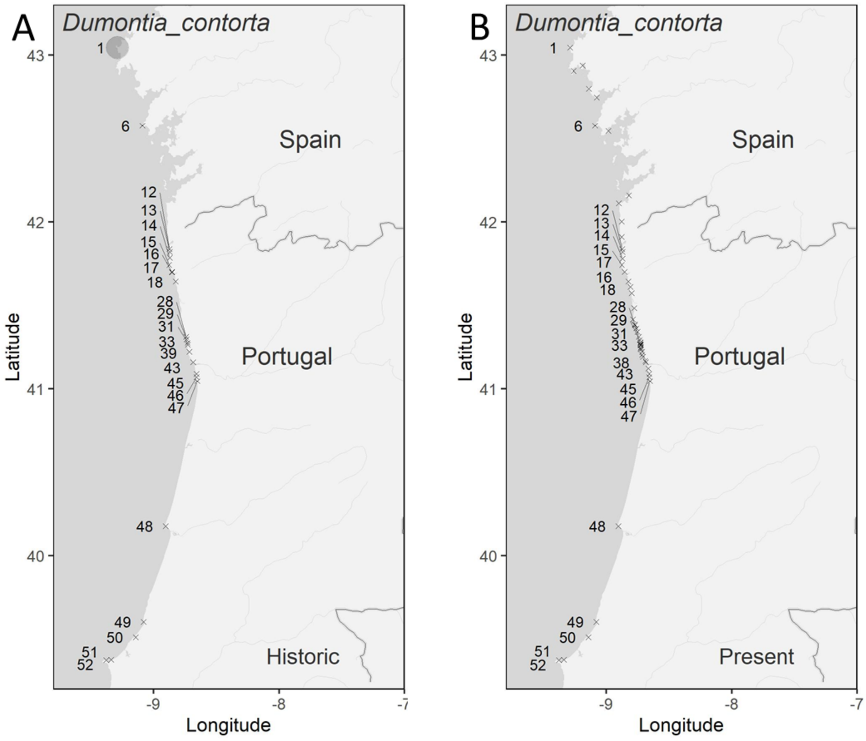

**Supplementary Figure 18** - Distribution maps of *Dumontia contorta* derived from historical (A) and present-day data (B).

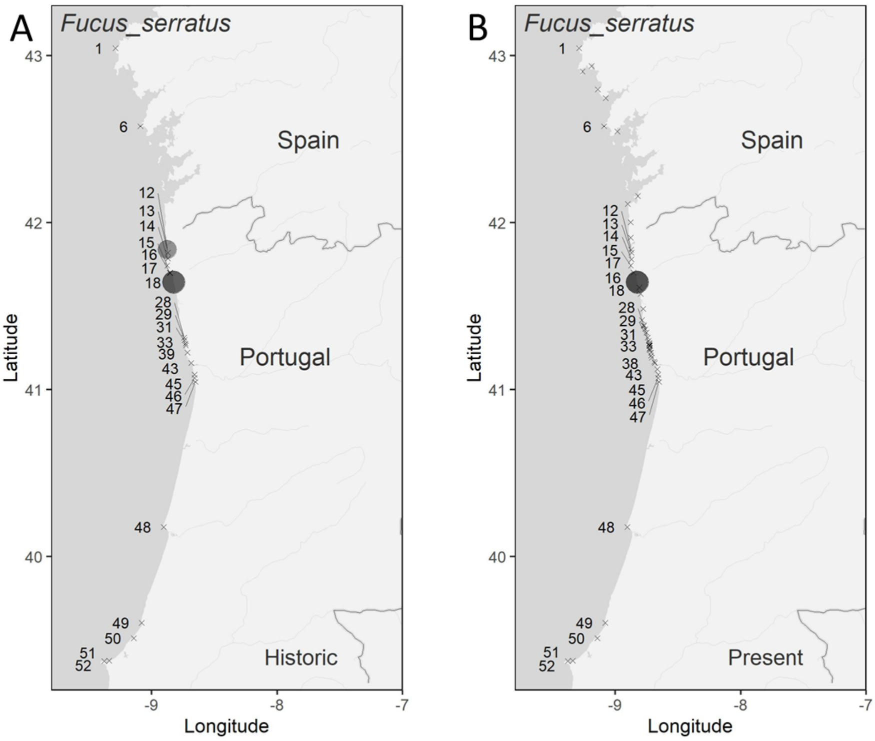

**Supplementary Figure 19** - Distribution maps of *Fucus serratus* derived from historical (A) and present-day data (B).

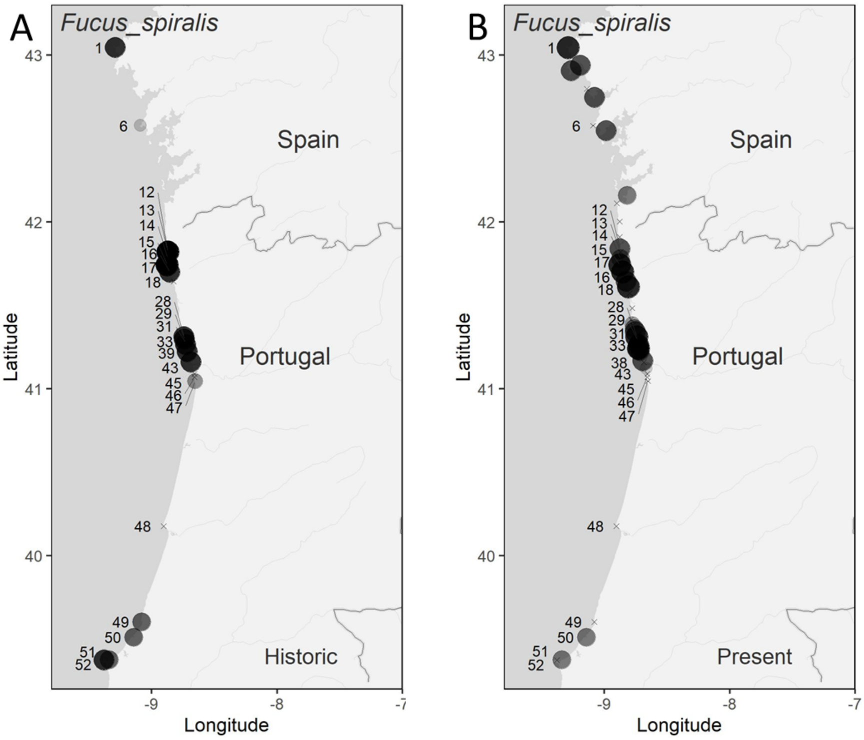

**Supplementary Figure 20** - Distribution maps of *Fucus spiralis* derived from historical (A) and present-day data (B).

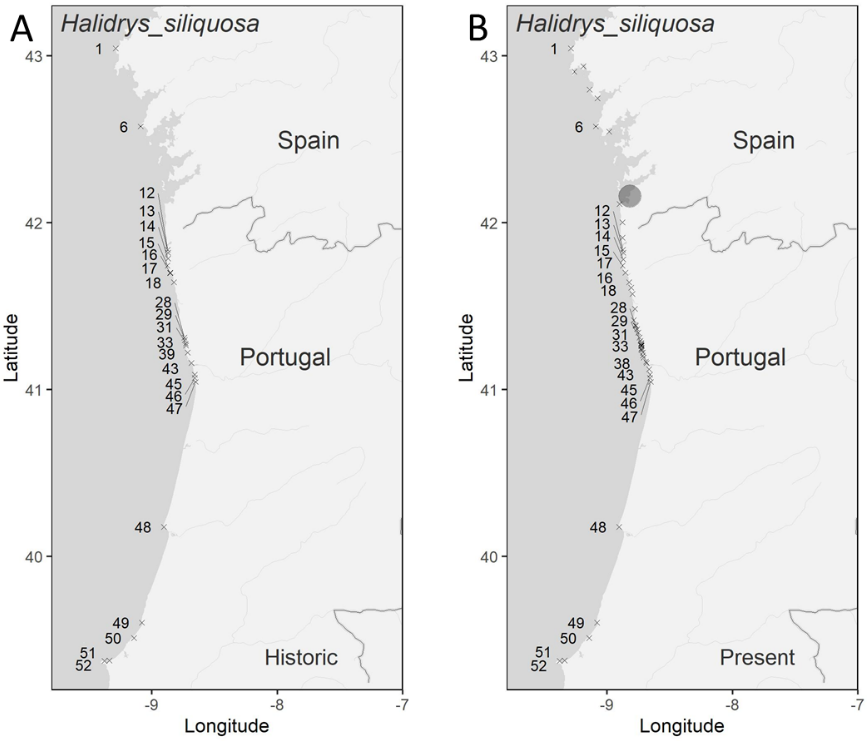

**Supplementary Figure 21** - Distribution maps of *Halidrys siliquosa* derived from historical (A) and present-day data (B).

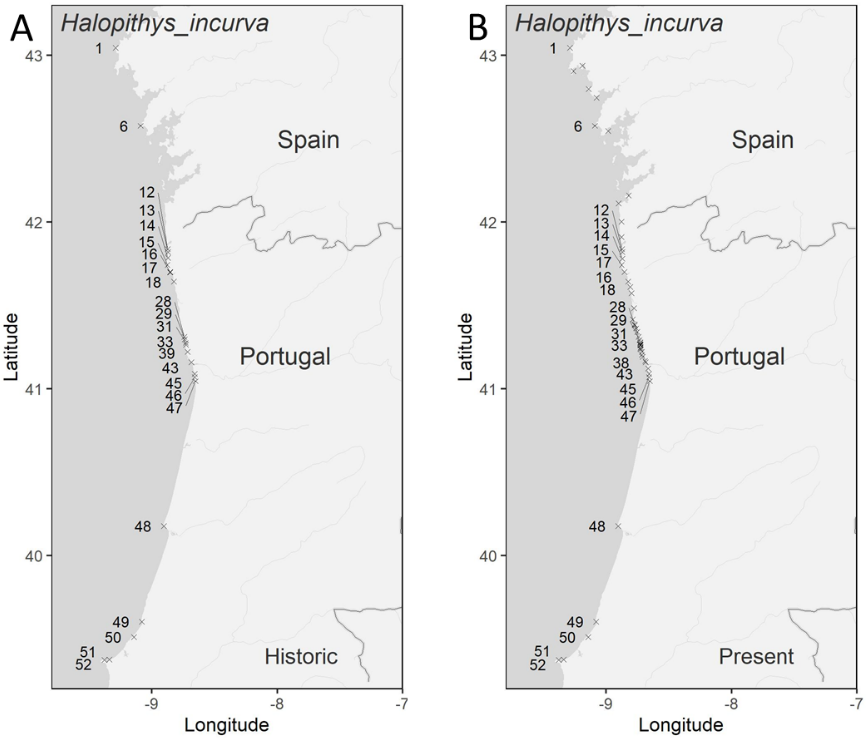

**Supplementary Figure 62** - Distribution maps of *Halopithys incurva* derived from historical (A) and present-day data (B).

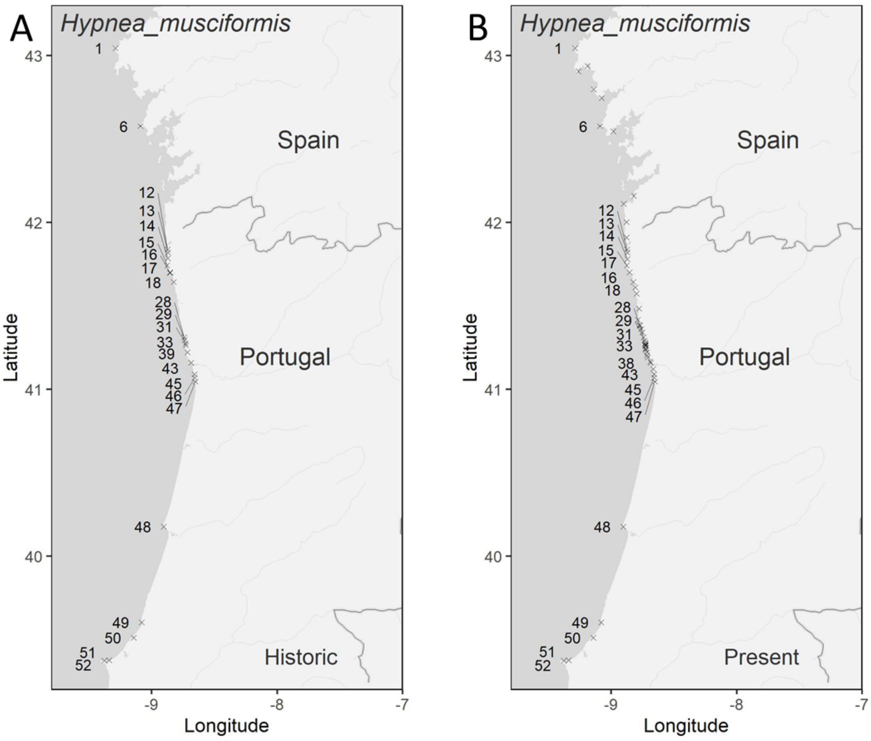

**Supplementary Figure 23** - Distribution maps of *Hypnea musciformis* derived from historical (A) and present-day data (B).

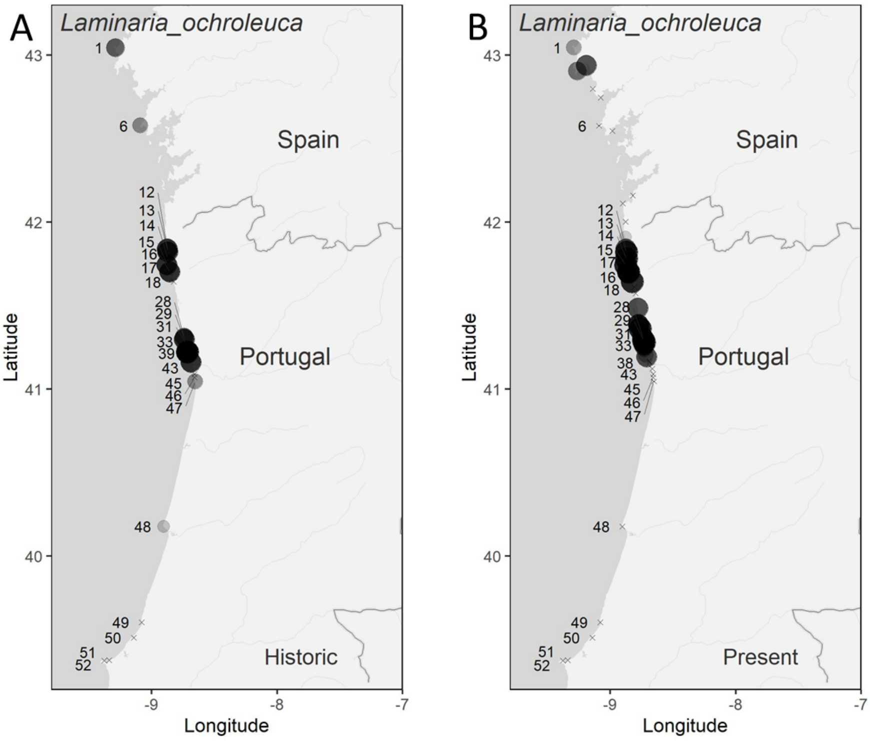

**Supplementary Figure 7** - Distribution maps of *Laminaria ochroleuca* derived from historical (A) and present-day data (B).

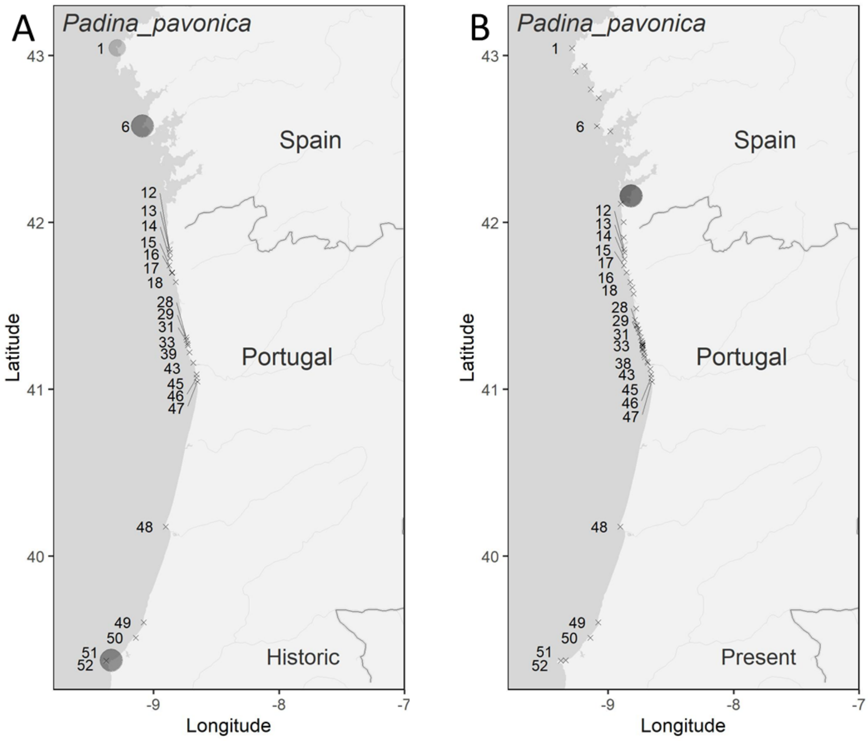

**Supplementary Figure 25** - Distribution maps of *Padina pavonica* derived from historical (A) and present-day data (B).

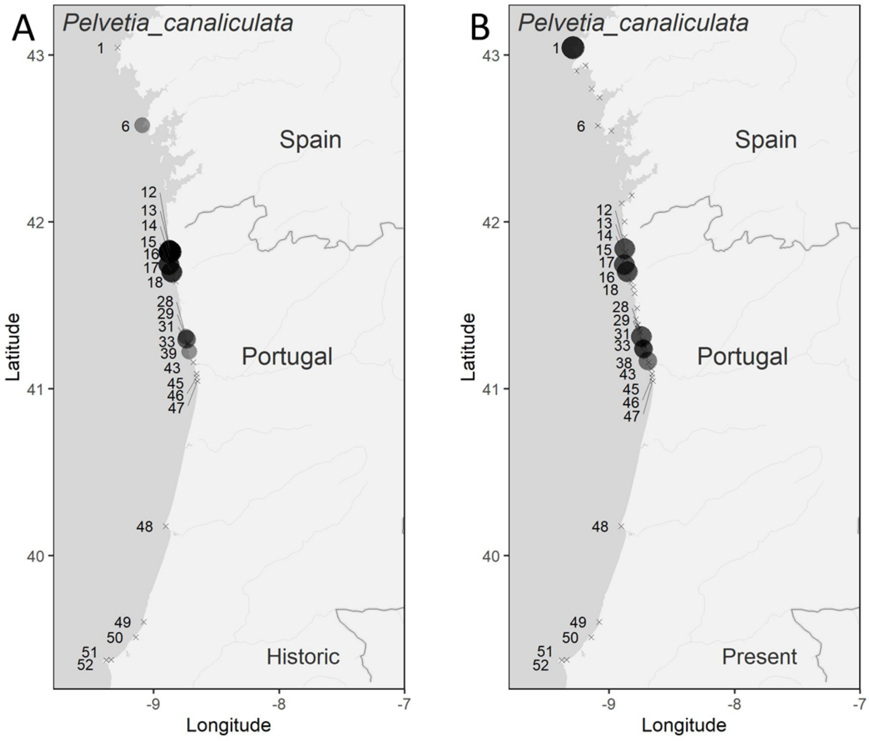

**Supplementary Figure 26** - Distribution maps of *Pelvetia canaliculata* derived from historical (A) and present-day data (B).

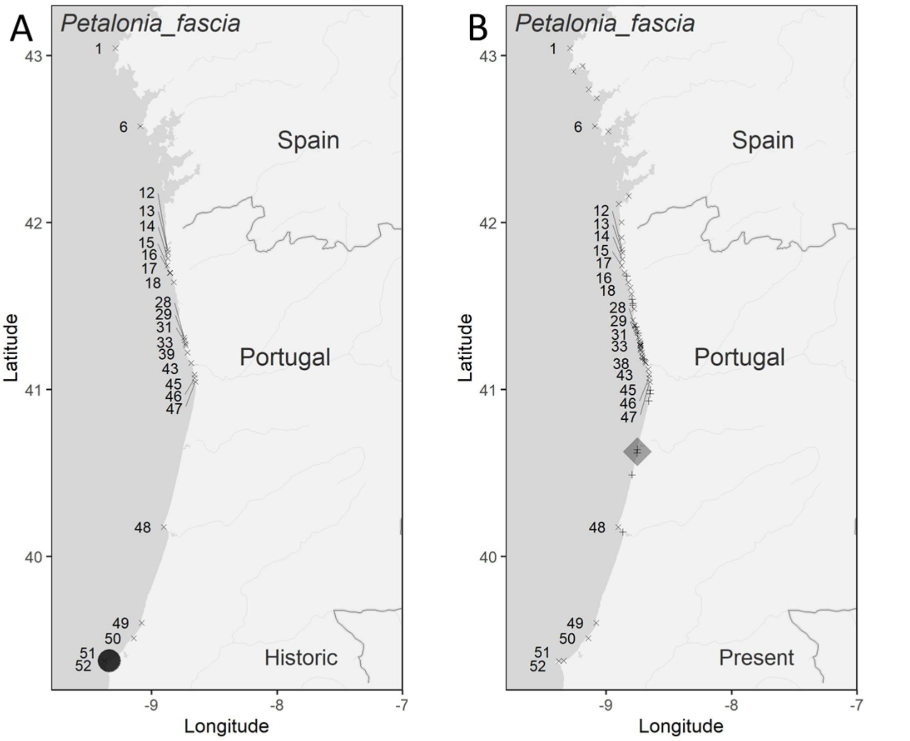

**Supplementary Figure 8** - Distribution maps of *Petalonia fascia* derived from historical (A) and present-day data (B). Rhombuses indicate the presence of the species in artificial sites and + indicate artificial sites in which the species was absent.

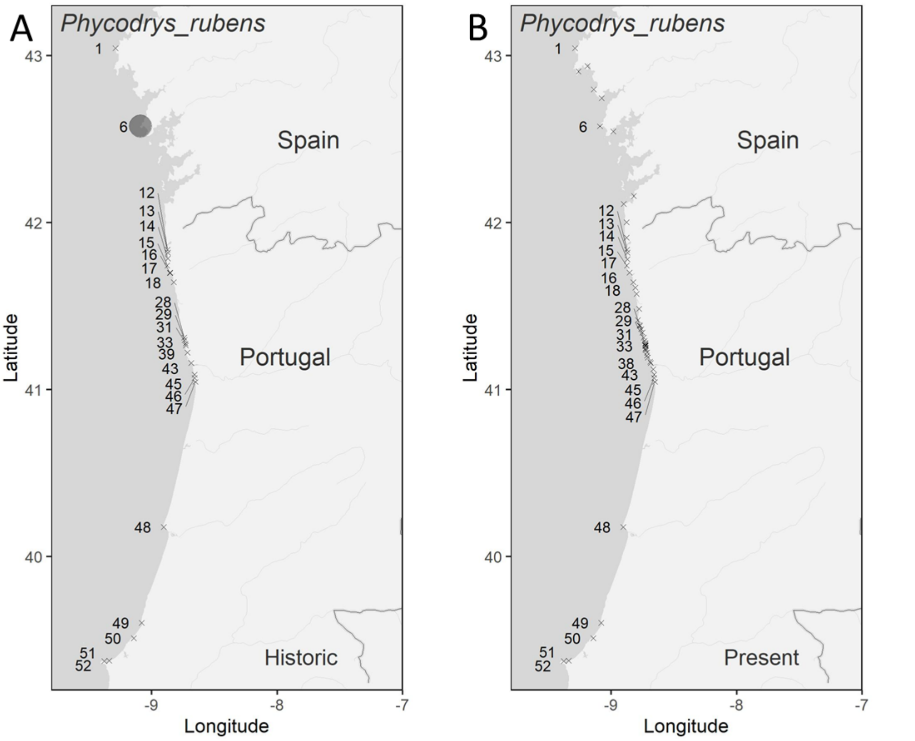

**Supplementary Figure 9** - Distribution maps of *Phycodrys rubens* derived from historical (A) and present-day data (B).

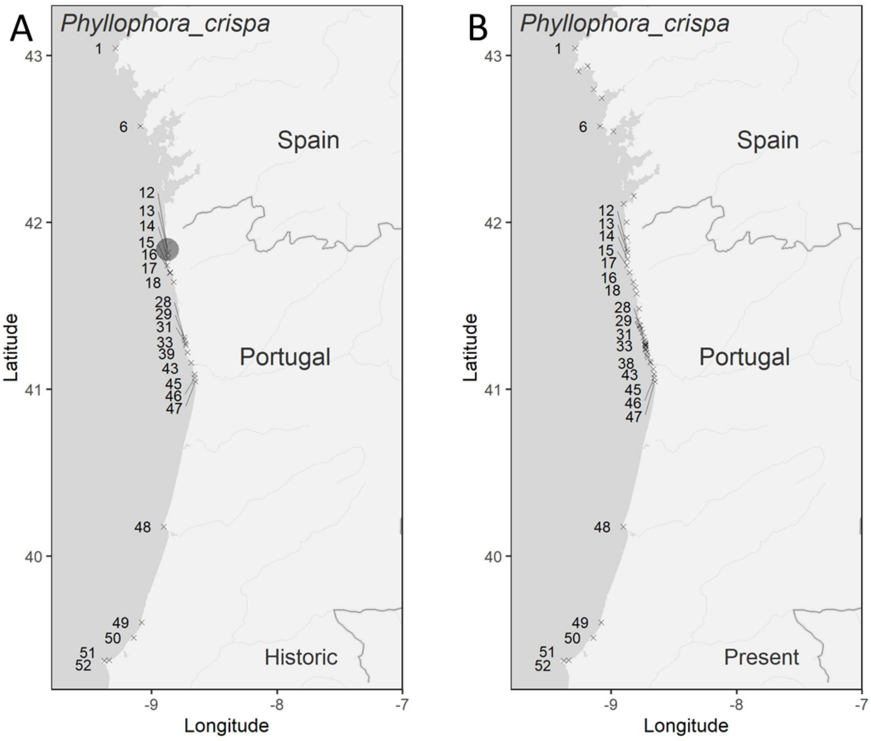

**Supplementary Figure 29** - Distribution maps of *Phyllophora crispa* derived from historical (A) and present-day data (B).

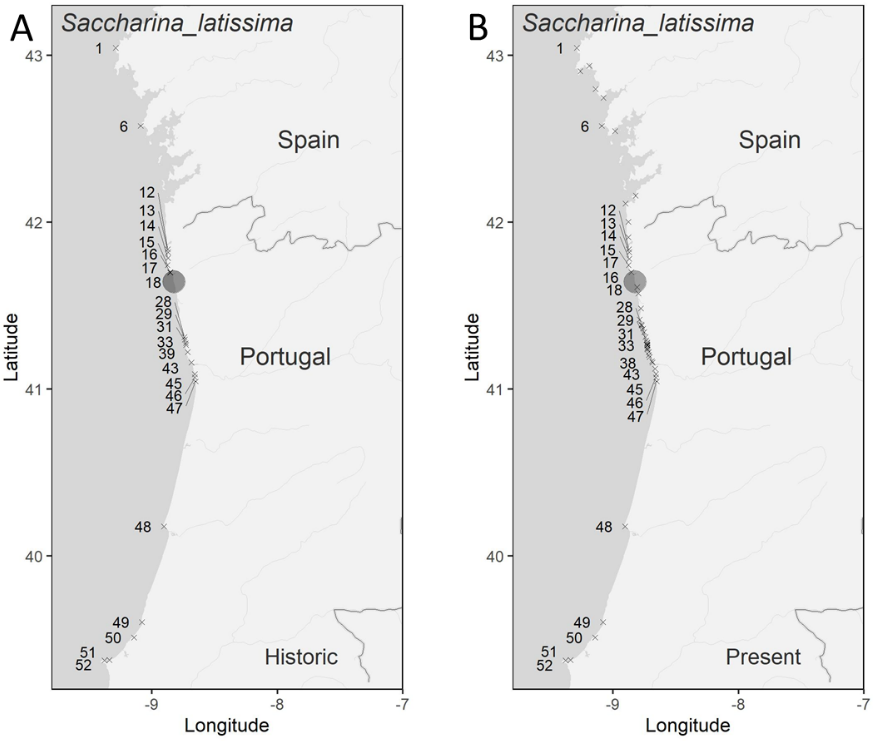

**Supplementary Figure 30** - Distribution maps of *Saccharina latissima* derived from historical (A) and present-day data (B).

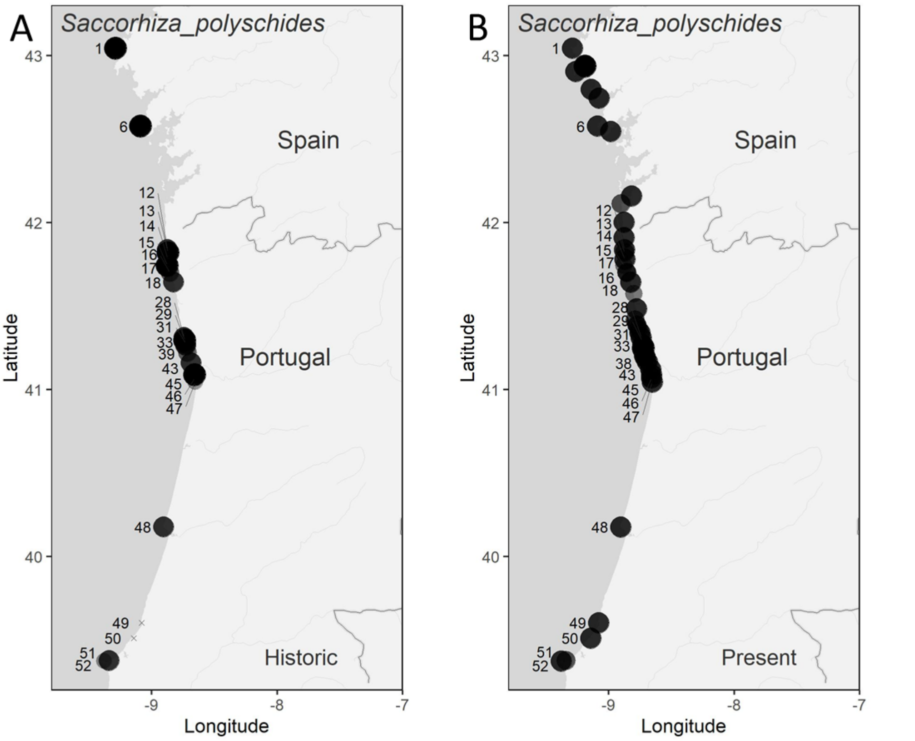

**Supplementary Figure 10** - Distribution maps of *Sacchoriza polyschides* derived from historical (A) and present-day data (B).

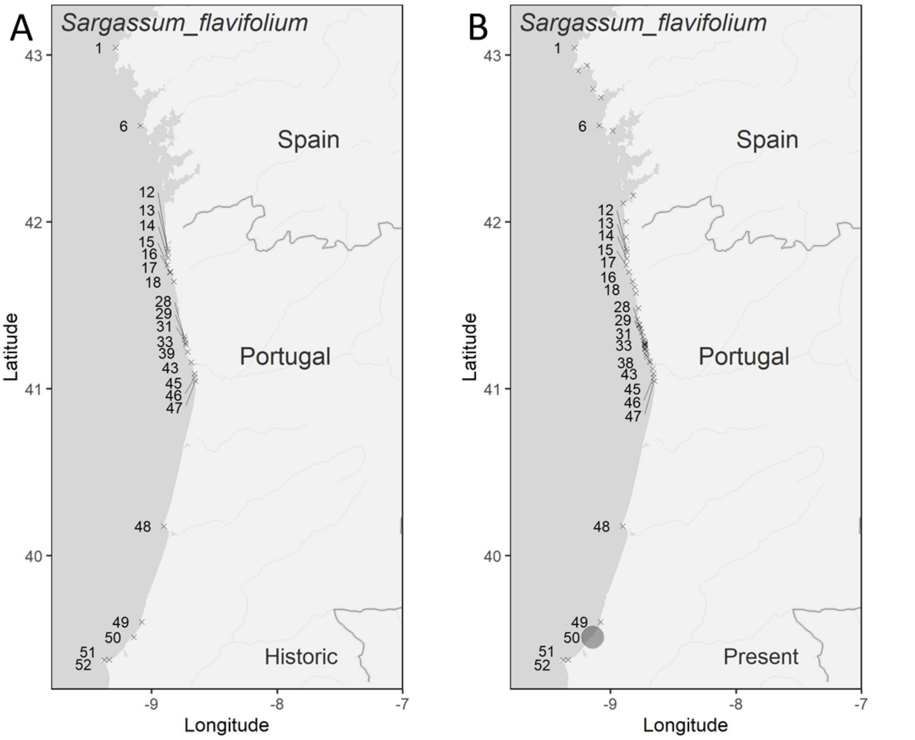

**Supplementary Figure 112** - Distribution maps of *Sargassum flavifolium* derived from historical (A) and present-day data (B).

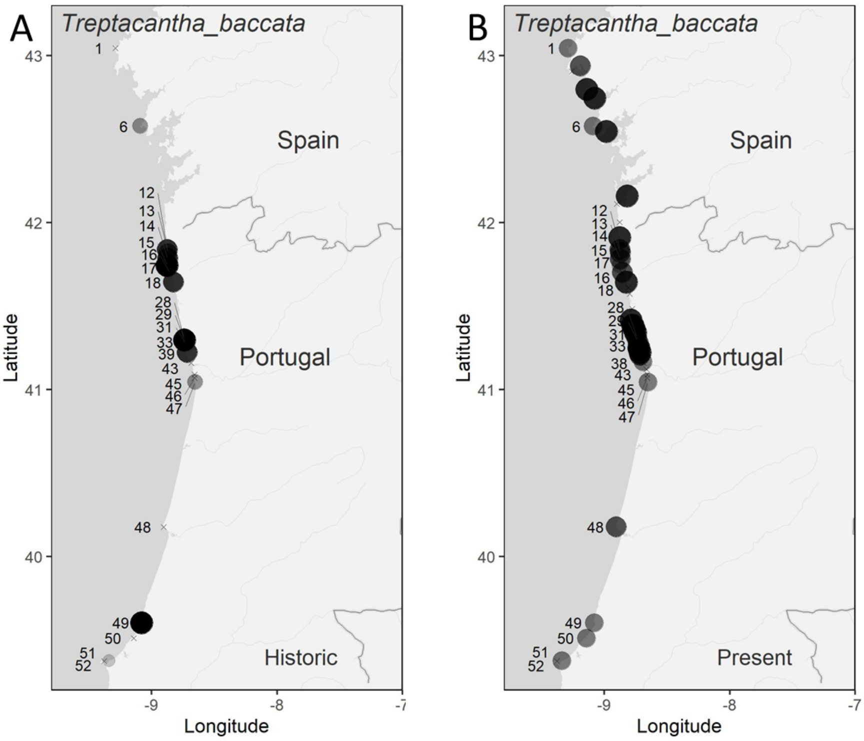

**Supplementary Figure 123** - Distribution maps of *Treptacantha baccata* derived from historical (A) and present-day data (B).

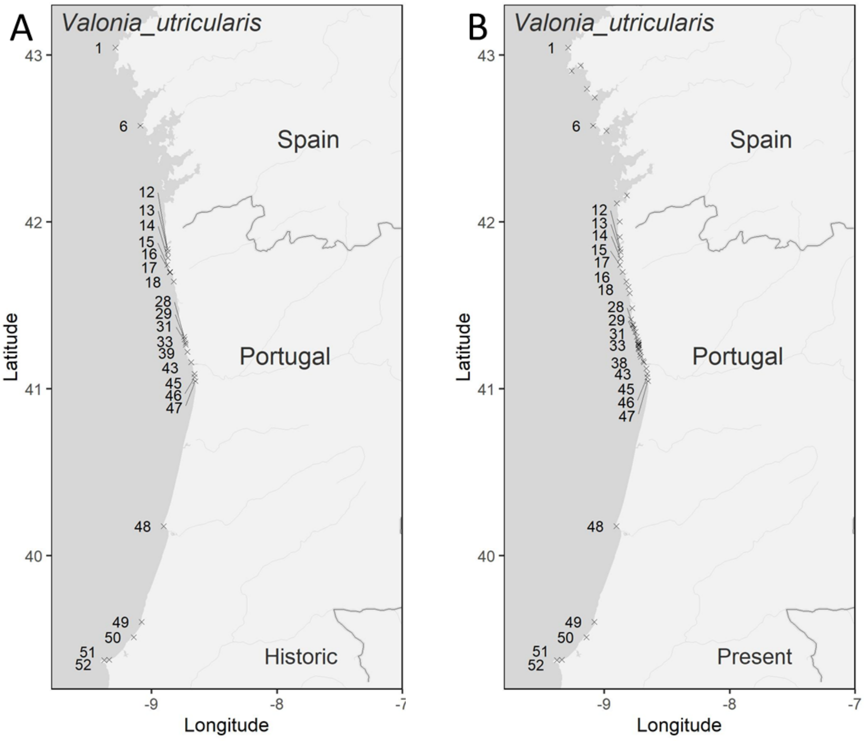

**Supplementary Figure 134** - Distribution maps of *Valonia utricularis* derived from historical (A) and present-day data (B).
